# Anterior Cingulate Cortex Directs Exploration of Alternative Strategies

**DOI:** 10.1101/2020.05.23.098822

**Authors:** D. Gowanlock R. Tervo, Elena Kuleshova, Maxim Manakov, Mikhail Proskurin, Mattias Karlsson, Andy Lustig, Reza Behnam, Alla Y. Karpova

**Affiliations:** Janelia Research Campus, Howard Hughes Medical Institute, Ashburn, VA, USA; Institute of Higher Nervous Activity and Neurophysiology of the Russian Academy of Sciences, Moscow, Russia; Department of Neuroscience, Johns Hopkins University Medical School, Baltimore, MD, USA; SpikeGadgets, San Francisco, CA, USA

## Abstract

The ability to adjust one’s behavioral strategy in complex environments is at the core of cognition. Doing so efficiently requires monitoring the reliability of the ongoing strategy and switching away from it to evaluate alternatives when appropriate. Studies in humans and non-human primates have uncovered signals in the anterior cingulate cortex (ACC) that track the pressure to switch away from the ongoing strategy, and others that relate to the pursuit of alternatives. However, whether these signals underlie computations that actually underpin strategy switching, or merely reflect tracking of related variables remains unclear. Here we provide causal evidence that rodent ACC actively arbitrates between persisting with ongoing behavioral choice and switching away temporarily to re-evaluate alternatives. Furthermore, by individually perturbing distinct output pathways, we establish that the two associated computations – whether to switch away from the current choice, and the pursuit of alternatives – are segregated within ACC micro-circuitry.

---

Making good choices in uncertain and dynamic situations is a core challenge of intelligent behavior. The requisite flexible action planning, which is thought to be implemented by the prefrontal cortex, effectively maps environmental and internal states onto behavioral strategies and prospectively searches through this mapping to select a suitable strategy. In open-ended natural settings, however, this mapping is potentially infinite, making a search for the optimal strategy computationally intractable. An emerging view is that our brains meet this challenge by employing heuristic approximations of optimal solutions, arbitrating among a small set of concurrently monitored behavioral strategies before pursuing the formation of new ones (Cleland et al., 2001; Donoso et al., 2014). Defining the circuit logic that permits balancing pursuit of the ongoing strategy with exploration of monitored alternatives is therefore a central issue in the neural control of behavioral adaptation.

One prefrontal cortical area, the anterior cingulate cortex (ACC), has long been viewed as key to behavioral adaptation in complex settings, especially when task or temporal context discovered through experience and deliberation calls for overriding of the ongoing behavioral strategy (Shenhav et al., 2017). Indeed, functional imaging studies in humans have uncovered rich contextual representations of relevant evaluative signals in ACC (Shenhav et al., 2016), as well as ACC activation during a departure from the ongoing strategy to re-explore previously learned alternatives (Blanchard and Gershman, 2018; Cleland et al., 2001). Furthermore, findings in humans (Behrens et al., 2007; Kolling et al., 2012; Kolling et al., 2014; McGuire et al., 2014; O’Reilly et al., 2013; Schuck et al., 2015), non-human primates (Blanchard and Hayden, 2014; Hayden et al., 2009; Procyk et al., 2000) and rodents (Karlsson et al., 2012; Ma et al., 2016; Powell and Redish, 2016), have implicated ACC in keeping track of alternative learned strategies, and in learning higher order statistics about the environment that can convey the need to switch away from the ongoing strategy (Kolling et al., 2014; Quilodran et al., 2008). Despite the mounting evidence that neural signatures related to the decision to switch away from the ongoing strategy and explore others abound in ACC, whether this area plays an active role in strategy switching, or merely monitors environmental context and behavioral performance, remains unclear.

In this study, we sought to directly probe ACC’s role in ongoing evaluation of strategies by transiently perturbing its activity at specific times during the decision-making process. Using targeted optogenetic manipulations of ACC activity in a temporally-structured rat foraging paradigm, we demonstrate how perturbing ACC activity at different epochs of the task – specifically, the point of commitment to an action dictated by a given strategy versus the point when the strategy is re-evaluated – leads to opposing effects on persistence with the ongoing strategy. Combined with an electrophysiological signature of strategy switching that we uncover in targeted electrophysiological recordings of ACC ensemble activity, these findings demonstrate that ACC plays an active role in the evaluation of the strategy space. They also directly expose its dual role in determining whether to persist with the ongoing strategy and, when switching away is deemed contextually appropriate, in guiding the exploration of alternatives. Using viral-mediated access to individual ACC output pathways, we then reveal the circuit structure supporting this dual role, demonstrating that it relies on anatomically distinct circuits. Overall, our findings argue that an opponent interaction within ACC micro-circuitry determines the balance between persisting with an ongoing strategy and switching to evaluate the alternatives.

## Results

### Behavioral framework that exposes strategy changes informed by inferred global task features

We trained rats on a behavioral paradigm, described previously (Karlsson et al., 2012), that captures the essential structure of sequential decision-making during foraging (Barack and Platt, 2017; Stephens et al., 2004) by presenting a series of accept-or-reject choices for two tone-cued options, each paired with distinct reward probabilities and presented in a randomly iterated manner (Figure 1A, Experimental Procedures). On each self-initiated trial within this framework, rats are presented with one of two auditory tones while at the central initiation port. Each tone indicates an encounter with a specific choice option (‘left’ or ‘right’) that – with a certain probability – provides a liquid sugar reward at the corresponding reward port after an imposed ‘travel time’, implemented as a requirement to perform several lever presses. Animals are free to either accept the option by proceeding to press the corresponding lever for a possible reward or reject it by re-initiating at the central port. This temporally structured sequential encounter process is thought to more closely resemble natural patch/prey encounters during foraging than the more common experimental setting where the animal must choose between simultaneously presented alternatives, and is thus more likely to expose core decision-making processes. In order to provide sufficient environmental uncertainty to prompt strategy switching, each behavioral session comprised several blocks of distinct reward probability pairs; reward probability associated with each option changed independently to ensure that sampling of both options is required to ascertain that a change has occurred. As we reported previously (Karlsson et al., 2012), after the unsignaled block transitions, animals typically entered a bout of undirected resampling of both choice options (Figure 1B, dashed box). Following these initial transition phases, expert animals often displayed a strong commitment to a single (‘preferred’) choice option (Figures 1B, S1, S2) by accepting nearly all trials for which the encountered option was the ‘preferred’ one and rejecting most of the trials for which the option encountered was the ‘non-preferred’ one.

**Figure 1.**
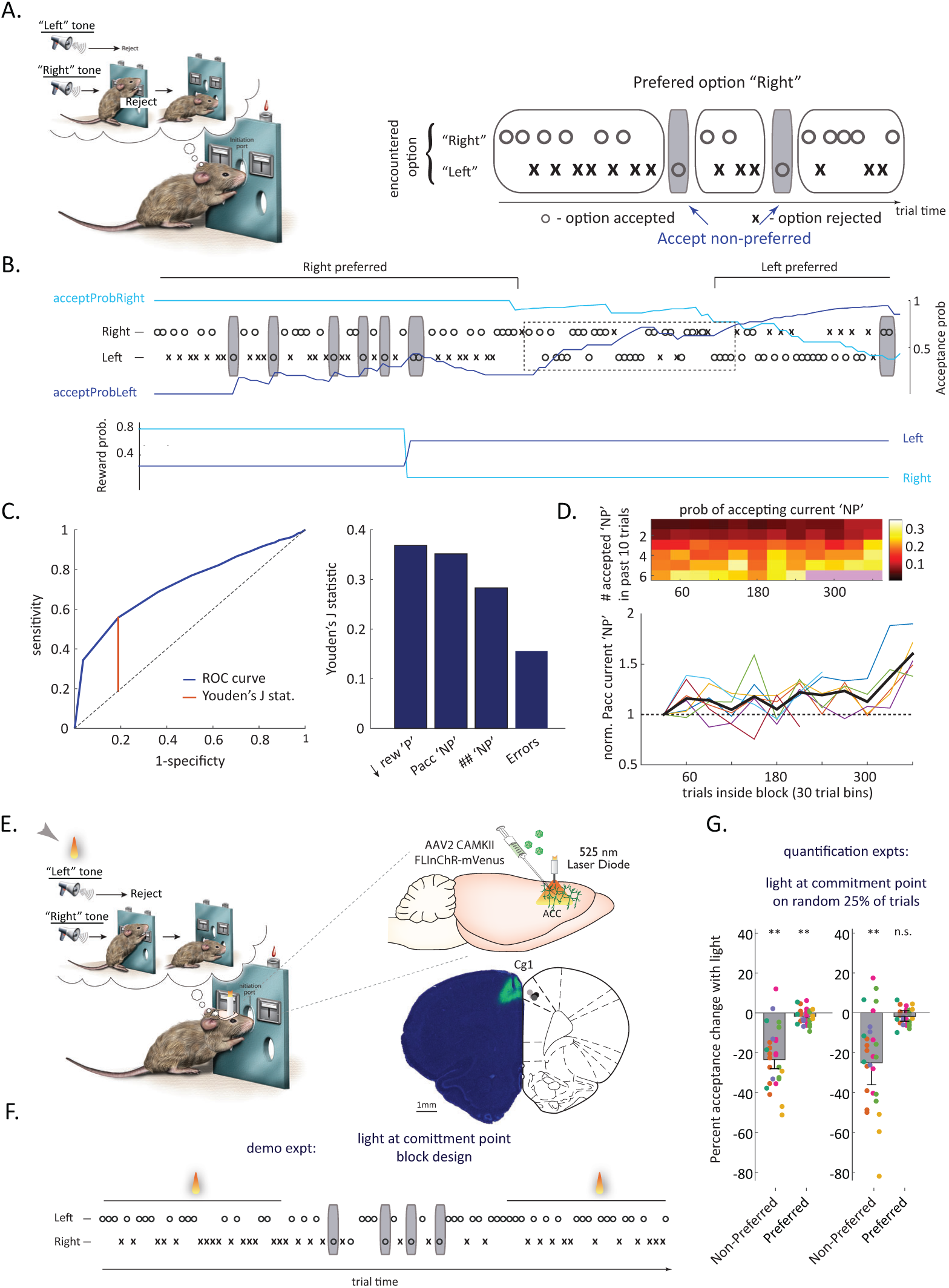
ACC perturbation at the commitment point leads to decreased pursuit of ‘non-preferred’ choice option in a foraging context. **A, Left panel:** Schematic of the task. Sequential, randomly iterated, tone-cued presentation of two behavioral options leads to a series of accept/reject decisions. **Right panel:** Block-wise structure promotes local resolve to pursue one option (here, ‘right’; also in the thought bubble in the task schematic) at the expense of the other (here, ‘left’). This default strategy tends to be occasionally interrupted by sporadic decisions to accept the non-preferred option (grey shading). **B, Top panel:** Sample behavioral trace around a block transition. Note a marked preference at the beginning (for ‘right’ choice option) and end (for ‘left’ choice option) of the behavioral trace, and the sporadic decisions to deviate from pursuing the ‘preferred’ option (gray shading). Dashed box highlights an example of previously described transient periods of resampling following block transitions. Solid blue and cyan line represent the independent measures of running acceptance probability for the ‘left’ and ‘right’ options respectively, used to identify ‘preferred’ and ‘non-preferred’ trials (see Experimental Procedures). **Bottom panel:** Block structure. Blue line - reward probability associated with the ‘right’ choice option; cyan-with the ‘left’. **C, Left panel:** Example ROC space that demonstrates the calculation of Youden’s J-statistic-a metric that captures the best dichotomization of data by a feature. Blue line – the ROC curve for a local downturn in reward of the ‘preferred’ side; dashed line – random classification; orange line – the maximum of Youden’s J-statistic (the difference between the ROC curve and the chance line that best separates the data into two classes). **Right panel:** Maximum Youden’s J-statistic for three behavioral features that display strongest discriminability: recent decrease in reward associated with the ‘preferred’ choice option (rew ‘P’), recent acceptance of the ‘non-preferred’ choice option (Pacc ‘NP’) and the number of consecutive presentations of the ‘non-preferred’ choice option (## ‘NP’). For comparison, Youden’s statistic was also calculated for one of the lesser predictive features, the local error rate (Error). **D, Upper panel:** Heat plot of the prevalence of accept decisions for the ‘non-preferred’ choice option on offer as a function of the number of trials inside a block of stable probabilities (x axis, values estimated for 30 trial bins) and of the number of acceptances of the ‘non-preferred’ choice option within the past 10 trials (y axis). Black/white color denote no/frequent acceptances of the ‘non-preferred’ option respectively; purple color denotes values without sufficient data. **Lower panel:** The same data normalized to the initial prevalence of the acceptance of the ‘non-preferred’ option. Each colored line represents the normalized prevalence of accept decisions for the ‘non-preferred’ choice option on offer given different parameterizations of the number of such accept decisions in the previous 10 trials. Black line represents the mean of these different parameterizations. **E, Left panel:** Schematic of the perturbation experiment. Light delivery (yellow light cone) was restricted to the period of tone presentation (grey arrow), when the animal commits to an action dictated by a specific strategy. **Right panel, top:** schematic of the viral approach for opsin delivery to the ACC and for local fiber implantation. **Bottom:** Opsin expression pattern. Circles on the atlas portion indicate locations of fiber placement. **F**, Accept/reject behavior in a demo perturbation experiment. Note greater acceptance of the non-preferred option (grey shading) during ‘no-light’ block compared with the two ‘light’ blocks. **G**, Illumination-dependent fractional change in the incidence of ‘accept’ decisions for the non-preferred and preferred options either throughout the behavioral session (left panel) or late into blocks of stable probabilities (right panel). n=6 animals. Different colors correspond to different experimental animals. Bars represent average effects across within-animal means. Error bars represent standard error of the mean. n.s.: not significant. **: p<0.01.

We next determined whether the animal’s choices in this task were guided simply by the local fluctuations in reward or were influenced by more complex task and temporal contexts, likely engaging the ACC. We were particularly interested in transient deviations from the ongoing strategy described above, when the pursuit of the ‘preferred’ choice option was occasionally interrupted by sporadic acceptances of the ‘non-preferred’ one (Figures 1A, B, gray rectangles). Such transient changes in strategy were observed deep into blocks of stable reward probabilities (Figure S3), and even when the difference in the payoff between the two options was large (Figure S4). This made a simple dependence on reward unlikely, and we turned to a quantitative analysis to describe the relationship between local task context and choice, and between more global temporal context and choice, thereby quantifying the influence of different factors. To permit the interpretations to be dissociated from the specific algorithmic approach the animals may be deploying, we used a Receiver Operator Curve (R.O.C.) analysis to examine how well the occurrence of particular behavioral features predicted animals’ choices when offered the ‘non-preferred’ option.

To increase the chances of capturing all the relevant behavioral information the R.O.C. analysis contained a large number of features. The task context-related features we examined fell into three main categories: the parameters associated with an individual trial leading up to the choice to be explained (at lags of 1-10 trials), running averages of different types of trials in recent past, and the number of trials since the last occurrence of a particular type of trial (see Experimental procedures for full list of features). We found that three behavioral features contributed prominently to the animals’ decision to accept the ‘non-preferred’ choice option: 1) recent (within the last 2-3 trials) reward omission associated with the ‘preferred’ option, 2) recent (within the last 10-15 trials) acceptance of the ‘non-preferred’ option (consistent with these decisions occurring in bouts) and 3) the run length of consecutive ‘non-preferred’ option encounters (Fig. 1C and Experimental Procedures). The bout-like nature of the deviations from rejecting the ‘non-preferred’ option suggests that when it comes to outcome evaluation, a local downturn in experienced reward (lack of reward associated with the ‘preferred’ option on a given encounter) is not the only factor driving the observed strategy switching. Indeed, decisions to persist after such local downturns were not obviously driven by the ‘preferred’ option being otherwise rewarded more or less in the recent past (Figure S5, no significant difference in more than half of the sessions in the distributions of reward history between unrewarded ‘preferred’ trials that were followed by the decision to persist or switch, Kolmogorov-Smirnov test). Furthermore, the strong and reliable contribution to the ‘non-preferred’ acceptance events from local task-related features (Fig. 1C, right) argues that they are unlikely to be random, unintended failures to choose the ‘correct’ behavioral response (that is, a response aligned with the ongoing default strategy).

We next asked whether additional more global features affect the animals’ decision, focusing in particular on temporal context. We measured the residuals from the local-feature prediction model and consistently observed a gradual increase in the prevalence of acceptances of ‘non-preferred’ choice option as a function of time within a block (Figures 1D, S6). Thus, a variable that correlates with time within a block, such as a growing awareness that a block may be nearing its end that reflects the brain’s ability to estimate environmental volatility (Behrens et al., 2007; Nassar et al., 2010), also contributed to the decisions to accept the strongly ‘non-preferred’ option. Together, these analyses revealed that the animals did not simply follow fluctuations in local reward but integrated task and temporal context when choosing their strategy.

### Temporally structured decision making task allows perturbation at specific time points with different computational relevance

To begin to explore the role of ACC and its neural underpinnings, we optogenetically perturbed ACC activity at two critical and qualitatively different time points within a trial: 1) the period of tone presentation during which time the animal learns of the identity of the encountered option and commits to an action dictated by his chosen strategy, and 2) immediately after the delivery of feedback at trial’s end during which time the success of the chosen action or strategy is likely evaluated and the decision to either persist with the ongoing strategy or to explore an alternative is made. We ensured spatial selectivity of our perturbation by expressing a recently developed, robust light-dependent silencer FLInChR (Brown et al., 2018) locally within the ACC (focusing specifically on region 24b within Cg1 ((Vogt and Paxinos, 2014), Figure 1E and Experimental Procedures), and by using modest light intensities (∼ 1-2mW). Applying the perturbation at the time of tone presentation, we found a striking difference between its effect when the presented choice option was the ‘preferred’ one, i.e., one that in the recent behavioral past the rat preferentially accepted, and when it was the ‘non-preferred’ one (see Figure 1 and below). To simplify the analysis, we thus focused on periods where there was a sufficiently large relative preference for one side, thus making one option clearly ‘preferred’ (Experimental Procedures, such periods constituted about 75% of the dataset, Figure S2). FLInChR-expressing rats performed 700-1500 iterated accept-or-reject decisions (trials), a subset of which were paired with light delivery through bilaterally implanted optical fibers, with the duration of the light stimulus restricted to the period of tone presentation (Figure 1E, Experimental Procedures). This experimental design permitted us to compare behavior for ‘light-on’ vs ‘light-off’ trials within individual sessions, making the analysis robust to individual differences in the overall incidence of accept decisions for ‘non-preferred’ trials, which varied across sessions.

### ACC perturbation at the time of choice option encounter preferentially decreases acceptance of the ‘non-preferred’ option

Application of light pulses during tone presentation led to a marked systematic reduction in the incidence of accept decisions for ‘non-preferred’ trials, with over 2-fold suppression observed in some sessions (Figures 1F, G, ratio of accept decisions for ‘light-on’ vs ‘light-off’ trials 0.77 +/-0.05 across within-animal means p < 0.005, Wilcoxon rank sum, n=6 animals, see also Figure S7A for the effect on absolute acceptance probability). The effect of ACC silencing on the incidence of accept decisions for ‘non-preferred’ trials was equally strong when transition phases following block changes were excluded from the analysis (Figure 1G, ratio of accept decisions for ‘light-on’ vs ‘light-off’ trials 0.75 +/- 0.09, p < 0.005, Wilcoxon rank sum, n=6 animals). To eliminate the possibility that the difference in the distribution of accept decisions for ‘light-on’ and ‘light-off’ trials occurred by chance, we repeated our analysis after randomly shuffling the ‘light-on’ and ‘light-off’ labels within individual sessions (Figure S8A). If our observations arose by chance, such a manipulation would be expected to frequently produce values of mean fractional change in acceptance on the order of what we observed experimentally. Instead, the observed light-dependent mean fractional changes in the acceptance rate lay outside of the corresponding ‘null’ distribution (Figure S8A). Shuffle analysis also confirmed that the effect of ACC perturbation on the acceptance on the ‘non-preferred’ choice option was consistent across experimental animals (Figure S9A). In contrast, light application had a minor (<2%) effect on the incidence of accept decisions for ‘preferred’ trials (Figures 1F, G, ratio of accept decisions for ‘light-on’ vs ‘light-off’ trials 0.98+/-0.01 across within-animal means, Figures S7A, S8A) that was not consistent across individual animals (Figure S10A). ACC perturbation thus preferentially decreases acceptance of an option that is currently not the ‘preferred’ one.

Several experimental factors (expression efficiency, efficiency of light delivery and variation in the resulting dynamics within the local ACC circuitry) likely contributed to the observed session-to-session variability in the perturbation effect; the extent of suppression on ‘light-on’ trials did not appear to correlate with the baseline rate of acceptance (one on ‘light-off’ trials) for the ‘non-preferred’ option within individual behavioral sessions (Figure S11). The observed behavioral change did not result from a light-dependent increase in occasional (<5%) erroneous attempts to accept the option not indicated by the tone (ratio of errors for ‘light-on’ vs ‘light-off’ trials 0.88 +/- 0.07; error trials were not analyzed further and are omitted from the examples shown). Nor could it have stemmed from an inability to process the auditory tones or from light-dependent aversion to a specific side of the behavioral box because within-session changes in reward probabilities ensured that unaffected preferred and affected ‘non-preferred’ trials comprised both choice options (‘left’ trials, paired with tone 1, and ‘right’ trials, paired with tone 2). More parsimoniously, the observed behavioral change likely stems from the context (‘non-preferred’ trial) in which the accept-reject choice is made.

We next used targeted extracellular recordings of neural activity to verify that ACC itself plays an active role in the decision to accept a ‘non-preferred’ option rather than passively providing nonspecific excitatory drive to a hypothetical key downstream area in which this computation actually occurs (Otchy et al., 2015). We reasoned that if ACC is selectively engaged during the decision to accept a ‘non-preferred’ as opposed to the ‘preferred’ choice option, then the neural representation associated with a specific action (e.g. ‘accepted right’) likely differs depending on the strategy-related context (‘preferred’ or ‘non-preferred’) (Figure 2A). Indeed, a substantial fraction of units recorded in ACC exhibited a marked difference in activity between the two conditions specifically during presentation of the relevant tone, with some units exhibiting a significant increase and others a decrease in activity (significant modulation in 42% of all neurons active during trial initiation, Figure 2B and Experimental Procedures). In contrast, the ensemble representation in such different behavioral contexts remains stable even in the supplementary motor area (SMA) – a region functionally coupled with ACC for context-specific behavioral adaptation (Bonini et al., 2014) – suggesting that this shift in encoding does not represent a brain-wide modulation of activity (*Proskurin et al, in preparation*). The separation of ACC ensemble dynamics associated with acceptance of a choice option between phases when the trial was ‘preferred’ or ‘non-preferred’ was sufficient to permit training of a linear classifier (Figure 2C, classification accuracy 75.2 +/-1.2 %, n=4 animals, N=25 sessions), suggesting that the neural pattern specifically associated with this difference is distinct enough to be read out by downstream networks.

**Figure 2.**
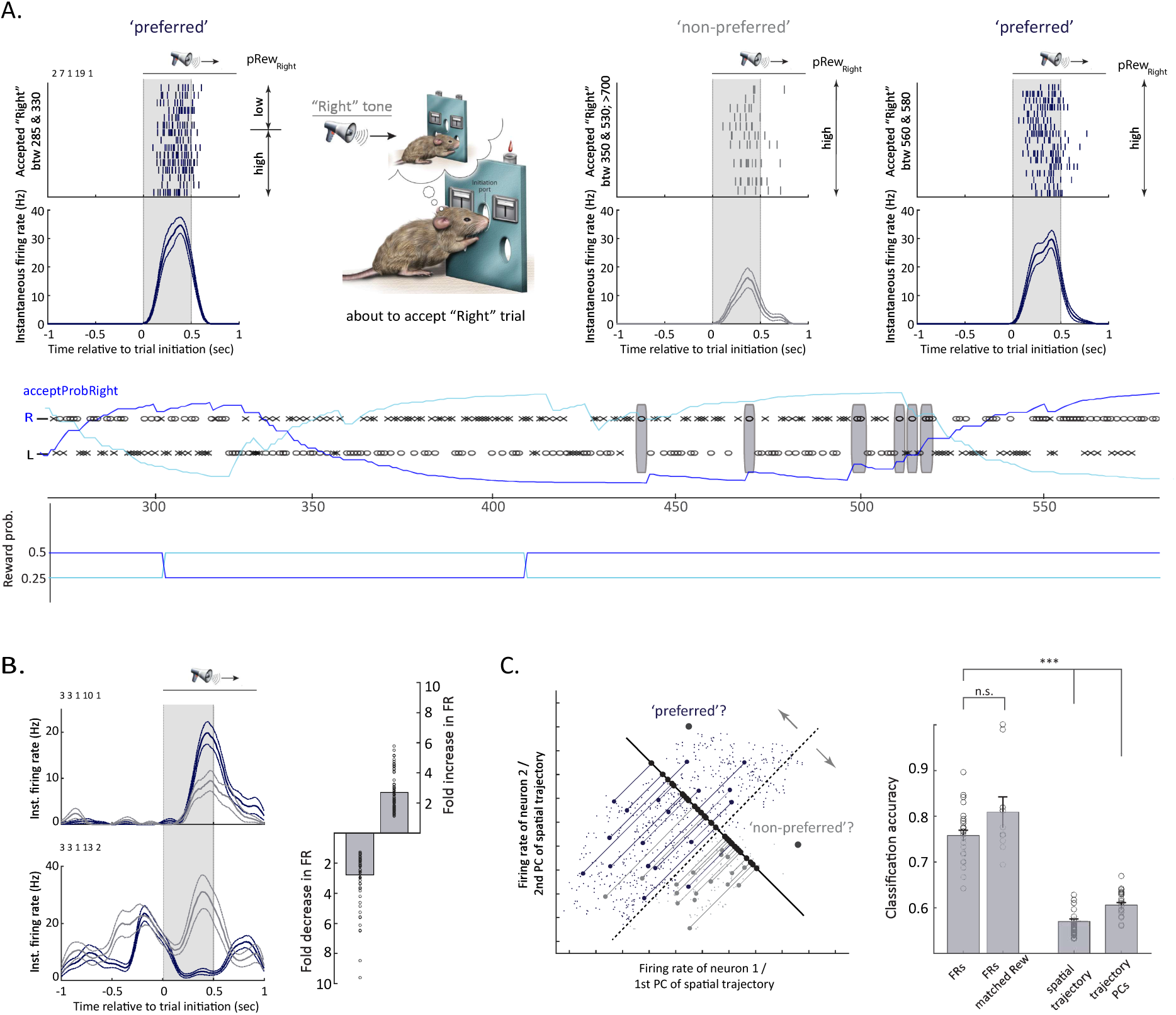
ACC ensemble activity reflects choice preference context. **A**, Top panel: Activity of an example cell around the point of commitment to an accept decision for the ‘right’ choice option, as that choice option was first ‘preferred’, then ‘non-preferred’ and back to ‘preferred’. Boxed region indicates the 500 msec window used for the decoding analysis in Figure2C. Middle panel: The corresponding behavioral trace. Gray rectangles indicate trials when the ‘right’ choice option was accepted despite being strongly ‘non-preferred’. Bottom panel: block structure. **B**, Left panel: Two simultaneously recorded cells that show opposite modulation by ‘preferred’ vs ‘non-preferred’ context. Blue: average activity on trials of the same type that are ‘preferred’; gray: ‘non-preferred’. Right panel: summary of fold modulation for all cells that show differential activity depending on the ‘preferred’ vs ‘non-preferred’ context. FR: firing rate. **C**, Left panel: simplified schematic of the linear classification (decoding) approach. Test data is projected onto the hyperplane that gives the best separation of the training dataset. Classification is done by comparing that projection to the threshold determined on the training dataset. Right panel: average classification accuracy (bars). Each point represents trial classification accuracy for one session. FRs: classification accuracy for trials in the ‘preferred’ and ‘non-preferred’ groups based on ensemble firing rates. n=4 animals, N=25 sessions. FRs, matched reward: classification accuracy based on ensemble firing rates for trials subsampled from the two groups to match the reward contingencies. n=4 animals, N=10 sessions with block structure confined to probability reversals. Spatial trajectory: classification accuracy for trials in the two groups based on the associated spatial trajectory. 2-dimensional trajectories were binned independently in x and y over 100 msec, and represented as points in a 10-dimensional space. n=4 animals, N=25 sessions. Trajectory PCs: classification accuracy for trials in the two groups based on the three main principal components of the associated spatial trajectories. n=4 animals, N=25 sessions. Error bars represent standard error of the mean. n.s.: not significant. ***: p<0.001.

Trajectory information is represented prominently in the ACC ensemble activity (Remondes and Wilson, 2013)), and thus the comparison of ACC ensemble activity at the commitment point between strategy-related contexts is confounded by the fact that our animals would have often approached the initiation port using different trajectories originating at spatially distinct locations. To verify that the observed modulation of ACC ensemble activity does not simply reflect a difference in the underlying trajectories and the associated kinematics of specific actions executed as a part of the ‘preferred’ versus the ‘non-preferred’ choice, we repeated the linear classification analysis now focusing on spatial behavioral trajectories. Indeed, the classifier performed significantly worse when trained either on the entire trajectory, or on its most variable component (Figure 2C, Experimental Procedures), suggesting that modulation of ACC ensemble activity cannot be explained solely by gross motor confounds.

We also evaluated the possibility that the observed ensemble activity modulation merely reflects different reward contingencies. For this analysis, one would ideally compare ensemble activity for trials in either the ‘preferred’ or the ‘non-preferred’ groups, subsampled to match in local reward history. However, we do not know how individual animals integrate local reward history, especially over randomly interleaved presentations of the two options. We thus focused on a subset of sessions where the probabilities of reward assigned to each block had a simpler structure: reward probabilities were confined to simple reversals between two values (Figure 2A), thus permitting a classification of trials into either a ‘high reward’ or a ‘low reward’ category (Figure 2A and Methods). The switches in reward blocks were uncued, and animals typically continued choosing their previously ‘preferred’ option for some time, despite it being rewarded less than the alternative (e.g., in Figure 2A, the ‘right’ side reverts to being the lower rewarded one at ∼trial 300, but the behavior adapts around Trial 350, leaving half of the accepted ‘right’ trials in the first ‘default’ block in the ‘low reward’ category). Trials associated with the acceptance of a specific choice option in the ‘preferred’ and the ‘non-preferred’ groups could then be sub-sampled to match in the reward category. If the observed modulation of ensemble dynamics were determined by reward, we would expect a decoder of the preference of the animal to fail when trained and tested only on reward category-matched subsets. In contrast, we found that decoders did no worse when trained on reward category-matched datasets than on the non-reward matched datasets (including the entire dataset, Figure 2C, classification accuracy 76.7 +/- 1.2 %, n=3 animals, N=10 sessions). Thus, more parsimoniously, the observed modulation stems from the context of the specific strategy (‘preferred’ vs ‘non-preferred’), in which acceptance of a specific choice option was made. Combined with the perturbation experiments, these observations argue that ACC actively directs resampling of ‘non-preferred’ choice options.

### ACC perturbation during strategy re-evaluation preferentially increases subsequent acceptance of the ‘non-preferred’ option

As re-evaluation of an ongoing strategy of pursuing the ‘preferred’ choice option is most likely to occur at the end of a ‘preferred’ trial (Holroyd and Coles, 2002; Kolling et al., 2014) – after the delivery of feedback in the form of present, or, more likely, absent, reward – we tested the effect of ACC perturbation when it specifically targeted this period (Figure 3A). Application of light pulses following the completion of ‘preferred’ trials resulted in the opposite effect from that observed at the commitment point, markedly enhancing the probability of acceptance of the ‘non-preferred’ option encountered on the following trial (Figures 3B, C, ratio of accept decisions for ‘non-preferred’ trial following ‘light-on’ vs ‘light-off’ preferred trials across within-animal means 1.67 +/- 0.21, p< 0.01, Wilcoxon rank sum, n=5 animals, see also Figure S12A for the effect on the absolute acceptance probability). Shuffle analysis again confirmed that that the observed difference in the distribution of accept decisions for ‘light-on’ and ‘light-off’ trials could not have arisen by chance (Figure S13A), and that the effect of perturbation was consistent across individual animals (Figure S14A). Crucially though, light application had no effect on acceptance of the following ‘preferred’ trials (Figures 3B, C, ratio of accept decisions for preferred trial following ‘light-on’ vs ‘light-off’ ‘preferred’ trials 1.02 +/- 0.01 across within-animal means p>0.5, Wilcoxon rank sum, n=5 animals, Figure S12A). Thus, ACC perturbation at the time of presumed strategy re-evaluation selectively increases subsequent acceptance of the ‘non-preferred’ choice option.

**Figure 3.**
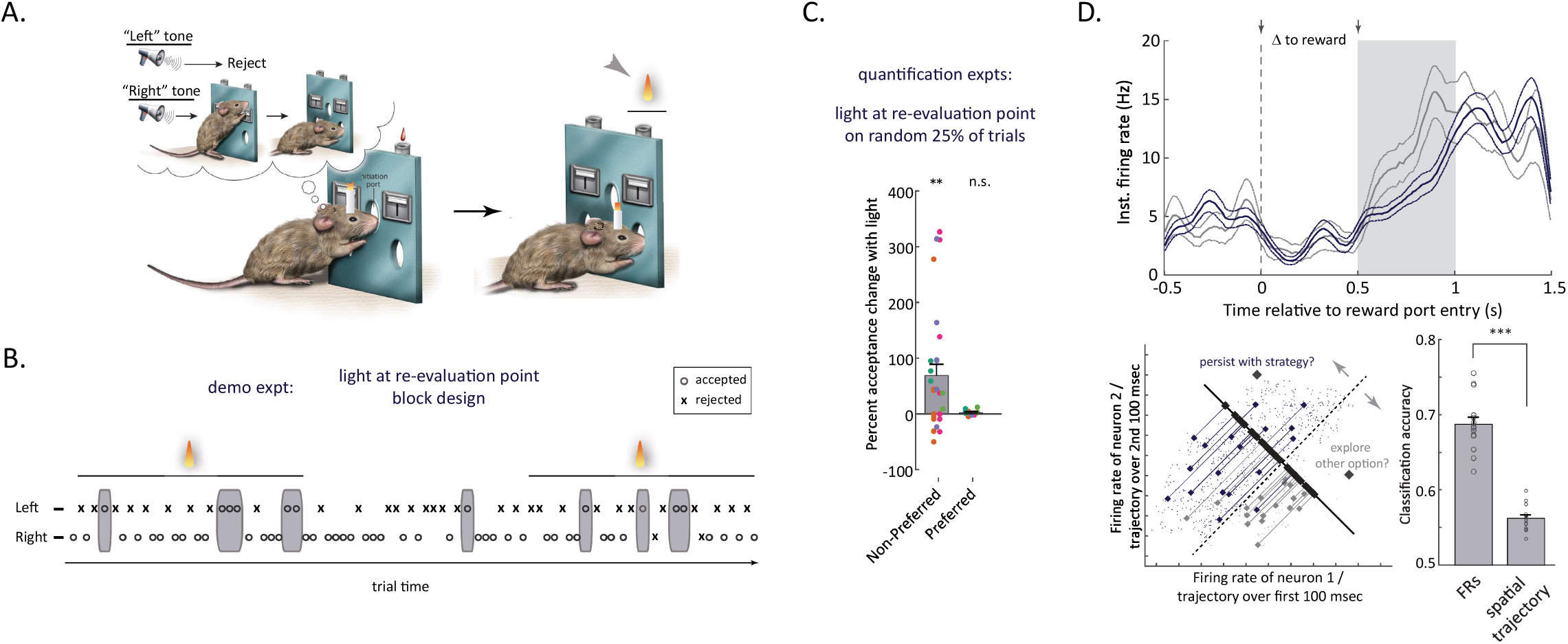
ACC perturbation during strategy re-evaluation leads to increased subsequent puruit of ‘non-preferred’ choice option. **A**, Schematic of the perturbation experiment. Light delivery (yellow light cone) was restricted to the feedback/re-evaluation point (gray arrow). **B**, Accept/reject behavior in a demo experiment. Note higher acceptance of the locally non-preferred option (black arrowheads) during ‘light’ blocks compared with the ‘no-light’ block. **C**, ACC illumination-dependent fractional change in the incidence of ‘accept’ decisions for the ‘non-preferred’ and ‘preferred’ options. Different colors correspond to different experimental animals. Bars represent average effects across within-animal means. n=6 animals Error bars represent standard error of the mean. n.s.: not significant. **: p<0.01. **D, Top panel:** An example cell that shows modulation by upcoming strategy switching during re-evaluation period after unrewarded preferred trials. Blue: average activity on trials when the animal decides to persist (i.e. reject ‘non-preferred’ option on next trial); gray: on trials before a strategy switch (i.e. before presented ‘non-preferred’ option was pursued). Boxed region from 500 msec to 1 sec indicates the window following the typical delay to reward delivery used for decoding. **Lower left panel:** simplified schematic of the linear classification (decoding) approach, as in Fig. 2. **Lower right panel:** average classification accuracy (bars). Each point represents classification accuracy for one session. FRs: classification accuracy for trials in the ‘persist with strategy’ and ‘explore the other option’ groups based on ensemble firing rates. Spatial trajectory: classification accuracy for trials in the two groups based on the associated spatial trajectory. n=3 animals, N=15 sessions. Error bars represent standard error of the mean. n.s.: not significant. **: p<0.01, ***: p<0.001.

To provide evidence in support of the conclusion that the observed perturbation effect indeed reflects the disruption of an active computation within ACC, we again examined the ACC neural ensemble dynamics. Specifically, we evaluated if, on the basis of ensemble activity, we could determine whether after a specific unrewarded preferred trial, the animal would persist with the ‘preferred’ strategy or switch. Although the markedly smaller sample size (of paired unrewarded preferred trials followed by non-preferred trials) posed challenges for statistical analyses, significant modulation of activity could still be observed at the level of individual cells (Figure 3D, upper panel); the majority (72%) of cells displaying significant modulation at the time of trial feedback did not show modulation at the commitment point. The observed modulation was sufficient to train a linear classifier that performed well on the abovementioned prediction task (Figure3D, lower panels, classification accuracy 68.6 +/- 1.0 %, n=3 animals, N=15 sessions). These findings argue that ACC also actively contributes to the computation that permits persistence in face of transient downturns and raises the question of whether these two opponent computations may be segregated by ACC micro-circuitry.

### The two perturbation effects can be anatomically dissociated via selective access to two ACC sub-circuits

A prominent role in behavioral persistence has been attributed to the intra-telecenphalic (IT) pathway and in particular its cortico-striatal branch (Langen et al., 2011), prompting us to evaluate if the cingulate’s contribution to the computation of whether or not to continue to persist with the ongoing strategy is likewise implemented within its IT sub-circuit. To gain independent access to the intra-telecenphalic pathway through its cortico-striatal (CS) component, we injected a retrograde viral vector (Tervo et al., 2016) that directed expression of FLInChR (AAV2-retro FLInChR-mVenus) bilaterally into the axonal terminal fields in dorsomedial striatum, and implanted optical fibers in the same location within the ACC as before (Figures 4A, S15A). The well-documented robust retrograde access to cortico-striatal cells (Tervo et al., 2016) permitted us to rely on direct delivery of the FLInChR transgene with AAV2-retro in the majority of the animals. However, to simplify subsequent comparison with the PT(CN) experimental group (see below), we also used a dual viral targeting strategy in a subset of the IT(CS) animals. This strategy combined a bilateral ACC injection of a regular viral vector that directed conditional expression of FLInChR (AAV-FLEX-FLInChR-mVenus) with a bilateral injection of a retrograde viral vector that directed expression of Cre recombinase (AAV2-retro Cre) into the axonal terminal fields in dorsomedial striatum (see Experimental Procedures). Having previously established that FLInChR permits robust pathway-specific perturbation of cortical circuits (Brown et al., 2018), we were able to examine the behavioral impact of perturbing the IT(CS) cingulate pathways by analyzing the influence of transient photoinhibition on the incidence of accept decisions for non-preferred trials at the two key time points within the task (commitment versus re-evaluation), as above.

**Figure 4.**
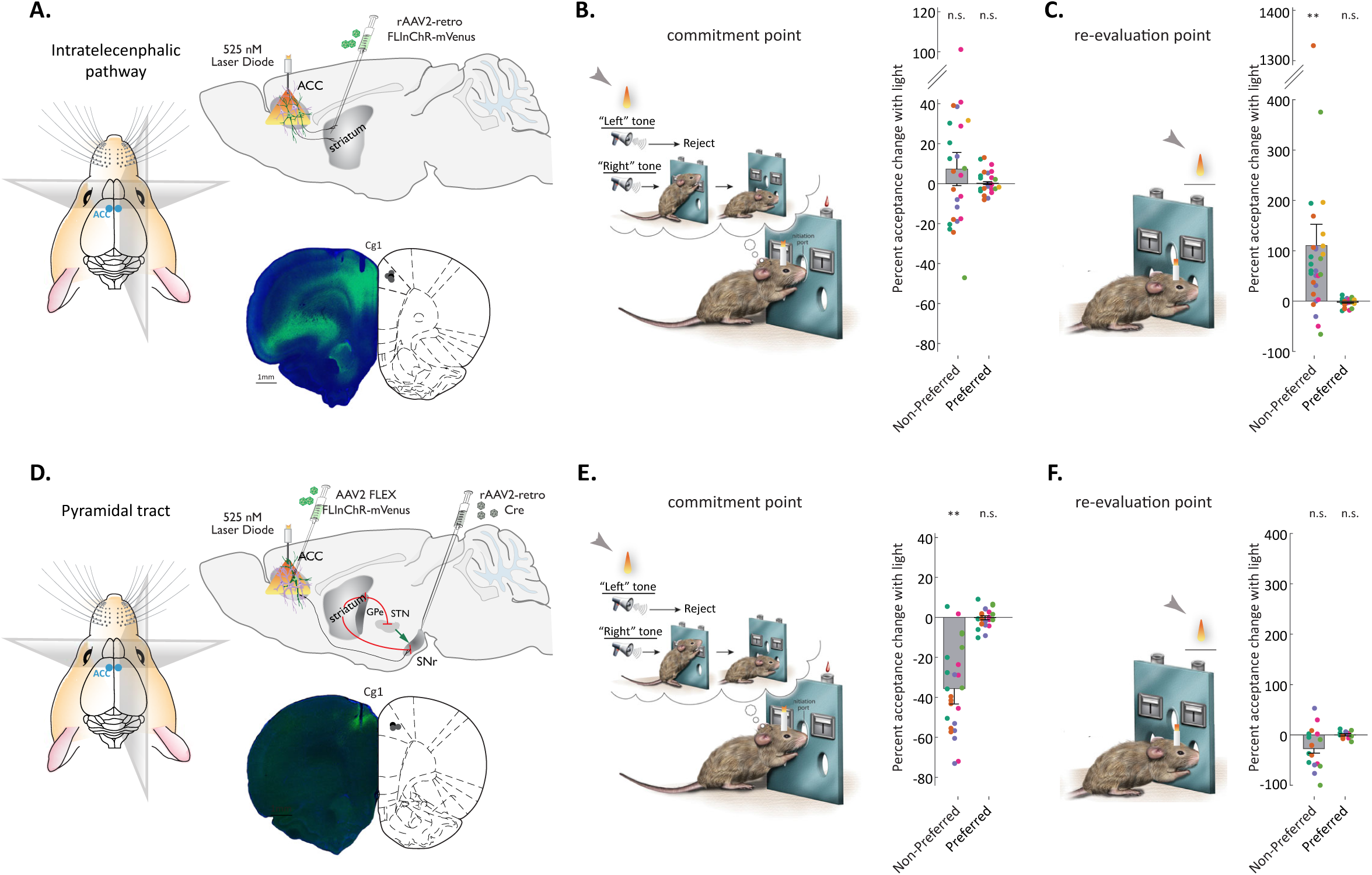
Computations associated with persistence and pursuit of ‘non-preferred’ choice option are segregated within the ACC micro-circuitry. **A**, Schematic of the viral approach for selective access to the intra-telecenphalic pathway through its cortico-striatal branch. Note that although opsin expression was not restricted to the IT(CS) pathway in one cortical region only, spatial specificity for ACC was preserved through localized light delivery. **B**, Fractional change in the incidence of ‘accept’ decisions for the ‘non-preferred’ and ‘preferred’ options as a result of cingulate’s IT(CS) perturbation at the commitment point. Note the lack of an effect on the acceptance of the ‘non-preferred’ option (cf. Figure1G). n=6 animals. **C**, Fractional change in the incidence of ‘accept’ decisions for the ‘non-preferred’ and ‘preferred’ options following cingulate’s IT(CS) perturbation at the time of default strategy re-evaluation (cf. Figure3C). n=6 animals. **D**, Schematic of the viral approach for selective access to the pyramidal tract through its cortcio-nigral (CN) branch. E, Fractional change in the incidence of ‘accept’ decisions for the ‘non-preferred’ and ‘preferred’ options as a result of PT(CN) perturbation at the commitment point (cf. Figure1G). n=5 animals. **F**, Fractional ch**ang**e in the incidence of ‘accept’ decisions for the ‘non-preferred’ and ‘preferred’ options following PT(CN) perturbation at the time of ongoing default strategy re-evaluation. Note the lack of an effect on the acceptance of the ‘non-preferred’ option (cf Figure3C). n=5 animals. Different colors correspond to different experimental animals. Bars represent average effects across within-animal means. Error bars represent standard error of the mean. n.s.: not significant. **: p<0.01.

Delivery of light pulses to the ACC of IT-FLInChR rats at the presumed point of re-evaluation resulted in a degradation of persistence that manifested in a marked increase of accept decisions for the following non-preferred trials, mirroring the effect observed when all of ACC was perturbed during the same task epoch (Figure 4C, ratio of accept decisions for non-preferred trial following ‘light-on’ vs ‘light-off’ preferred trials 2.10 +/- 0.42 across within animal means for all animals in the group, 2.32 +/- 0.43 when one mis-targeted animal was removed from the set, p< 0.01, Wilcoxon rank sum, n=6(5) animals; cf. 1.67 +/- 0.21 for all of ACC, Figure 3C, Figures S12B, S15A). Shuffle analysis again confirmed that that observed difference in the distribution of accept decisions for ‘light-on’ and ‘light-off’ trials could not have arisen by chance (Figure S13B), and that the effect of perturbation was consistent across all properly targeted animals, independent of the specific viral targeting strategy (Figure S14B). In contrast, the incidence of accept decisions for the ‘non-preferred’ option was not significantly reduced when perturbation in this group of animals was confined to the commitment point, thus not mirroring the effect of perturbing all of ACC during this task epoch (Figure 4B, ratio of accept decisions for ‘light-on’ vs ‘light-off’ trials 1.03 +/- 0.09 across within-animal means, p>0.05, Wilcoxon rank sum, n=4 animals; cf. decrease to 0.77 +/- 0.05 for all of ACC, Figure 1G; see also Figures S8B, S7B). These observations suggest that either the former computation is simply more sensitive to partial silencing of ACC activity, or that it is selectively implemented within the sub-circuit associated with the intra-telecenphalic pathway. The latter interpretation fits with the observed segregation of units that display activity modulation at re-evaluation or commitment points, and with the opposite behavioral effect of perturbing ACC during those epochs, but raises the question of which other ACC sub-circuit may implement the latter computation.

Recent work has highlighted the asymmetry of excitatory connectivity in the cortex, with intra-telecenphalic neurons impinging on subcortically-projecting pyramidal tract (PT) neurons (Brown and Hestrin, 2009; Morishima and Kawaguchi, 2006), providing, among other inputs, feedforward inhibition via local interneurons (Lee et al., 2014; Lu et al., 2017). These anatomical findings suggested that the PT pathway is a candidate downstream strategy switching circuit. We further reasoned that of the cingulate PT neurons, those projecting to substantia nigra pars reticulata (SNr) (Naito and Kita, 1994) – the output nucleus of the basal ganglia control – are potentially in a particularly privileged position to override ongoing behavioral strategies. The convergent input from the basal ganglia to the SNr, and the subsequent divergent outputs from the SNr to thalamus and superior colliculus, position the SNr as a bottleneck that may gate basal ganglia (BG) output. Indeed, the “direct pathway” striatal input and “indirect pathway” subthalamic nucleus input, converge onto a microcircuit in the substantia nigra that is believed to control the gain on this output (Brown et al., 2014; Gerfen and Paxinos, 2004). Thus, BG-dependent motor programs, such as switching away from an ongoing automatic behavior to explore an alternative option (e.g. Hikosaka and Isoda, 2008), could be profoundly impacted by a cortical input into the SNr (Doyon and Benali, 2005). ACC is a natural candidate controller of the SNr in such a model, given the multiple proposals that this cortical region contributes to switching away from an ongoing behavioral strategy. We thus sought to selectively targeted the cingulate PT pathway through its cortico-nigral (CN) branch for a similar set of perturbation experiments in a separate group of animals.

To gain independent access to the PT(CN) pathway, we again used a virally-mediated approach, combining a bilateral injection of a regular viral vector that directed conditional expression of FLInChR (AAV-FLEX-FLInChR-mVenus) into ACC with a bilateral injection of a retrograde viral vector that directed expression of Cre recombinase (AAV2-retro Cre) into the axonal terminal fields in the medial SNr (Figures 4D, S15B). Separately, we verified that bilateral striatal injections of AVV2-retro, which would be expected to target a subset of PT neurons that have axon collaterals in the ipsilateral striatum, do not efficiently target cotico-nigral neurons (Figure S16, double labeling observed in 1.24 +/- 0.79 % of cells). In FLInChR-expressing animals, we then similarly analyzed the influence of transient photoinhibition on the incidence of accept decisions for non-preferred trials, separately at the commitment point and at the point of re-evaluation. Strikingly, we observed a complementary pattern of behavioral impairment compared to that seen with IT-FLInChR animals. Indeed, perturbing the PT(CN) pathway now mirrored the effect on the incidence of acceptance decisions for the ‘non-preferred’ choice option observed following ACC perturbation at the commitment point (Figure 4E, ratio of accept decisions for ‘light-on’ vs ‘light-off’ trials decreased to 0.64 +/- 0.08 across within-animal means, p<0.01, Wilcoxon rank sum, n=5 animals, cf. decrease to 0.77 +/- 0.05 for all of ACC, Figure 1G; also see Figures S7C, S8C, S9B), but not at the time of strategy re-evaluation (Figure 4F, ratio of accept decisions for subsequent ‘non-preferred’ trial following ‘light-on’ vs ‘light-off’ ‘preferred’ trials decreased after PT(CN) perturbation to 0.73 +/- 0.09 across within-animal means, p<0.01, Wilcoxon rank sum, n=5 animals, cf. an increase to 1.67 +/- 0.21 for all of ACC, Figure 3C, and an increase to 2.32 +/- 0.43 for the IT(CS) pathway, Figure 4C; see also Figures S12C, 13C). Whether the unexpected *decrease* in acceptance of the subsequent ‘non-preferred’ option (as opposed to a lack of an effect) reflects bleed-through perturbation of the PT(CN) pathway as its activity slowly recovers following the offset of perturbation at that trial’s start, or some active process, the relevant local computation at this time point exposed by perturbing all of ACC or its IT(CS) pathway is unlikely to take place in this specific sub-circuit. Although these combined observations do not preclude a role for the IT and PT pathways in other aspects of adaptive behavior, they do indicate a degree of selectivity in the respective degradation of persistence and strategy switching. Overall, our findings provide strong causal evidence for ACC’s direct involvement in ongoing strategy arbitration in uncertain environments through a complex

## Discussion

Dissecting specific contributions that frontal cortices make to decision-making has remained challenging due to the experimental trade-off between using a model organism that is behaviorally sophisticated enough for flexible decision-making in complex and volatile environments, and one that permits robust circuit dissection with modern molecular tools. Here, we combine a foraging task in rats that requires animals to continuously arbitrate between strategies with a pathway-specific optogenetic perturbation to demonstrate that the anterior cingulate cortex (ACC) plays an active role in guiding strategy arbitration. Transient perturbation of ACC activity selectively during the period when a local downturn associated with the ‘preferred’ option presumably prompts strategy re-evaluation, resulted in an increased incidence of acceptance decisions for subsequent encounters of the ‘non-preferred’ choice option – a shift from the ongoing strategy of exploiting the ‘preferred’ one. Changing the timing of perturbation to the moment of commitment to an action dictated by a given strategy instead limited the pursuit of the available ‘non-preferred’ choice option. Targeting opsin expression selectively to, respectively, the IT and PT sub-circuits of the ACC permitted an anatomical dissociation of the two perturbation effects, and provided an internal control for any non-specific effect of viral infection or tissue illumination.

Our finding that perturbing ACC activity at the time of strategy re-evaluation does not lead to the ‘preferred’ option being rejected demonstrates that the ongoing strategy is not overridden unless the ‘non-preferred’ option is encountered, and argues that the decision of whether to switch away from the ongoing strategy and the actual pursuit of an action dictated by a specific alternative strategy represent two separate computations. Consistent with this interpretation, evidence for two independent, temporally-segregated signals – one indexing the growing pressure to pursue non-default choices and one related to the actual making of such choices once the pressure reaches a certain threshold – was found in a recent functional imaging study in human subjects (Kolling et al., 2014). Moreover, our pathway specific perturbation results suggest an intriguing possibility that there is an anatomical substrate to this functional duality within the ACC micro-circuitry: the output of the ACC’s PT sub-circuit drives pursuit of alternative learned strategies, but is kept clamped through local opponent interaction until evidence grows in favor of deviating from the ongoing strategy. If the representation that guides pursuit of alternative strategies is indeed embedded into a specific population of cortical projection neurons (the PT tract), then evolution was likely able to tap into one of its principle solutions for achieving a hierarchy of cortical processing: the dynamic segregation of discrete projection neuron ensembles via inhibitory control by specialized interneuron subpopulations. Indeed, the functional opponency between the IT(CS) and PT(CN) pathways – both excitatory – would necessarily involve an intervening directional inhibitory pathway.

What might be the nature of the computation subserved by the apparent functional opponency within the ACC micro-circuitry? One possibility is that it relates to the general tension between the need to persist with the ongoing strategy in face of transient downturns and the need to expeditiously detect, and adapt to – often unsignaled – changes in the environment that render past experience irrelevant for future predictions (Cohen et al., 2007). When such tension is informed by the temporal context as in the specific behavioral framework employed in our study, individual differences in resolving it may relate to differences in the estimation of environmental volatility, and a growing subjective estimate that a change is impending (Nassar et al., 2010). In this view, the presumably larger-scale network associated with the cingulate’s intra-telecenphalic pathway would play a key role in computations of the underlying internal temporal expectation. In turn, the unprompted switches away from the ongoing strategy would reflect release of the PT (CN) pathway from inhibitory influence once the subjective estimate of the impending change crosses a certain threshold. In other contexts, the relevant tension may be between making the default choice of persisting with the ongoing strategy and engaging cognitive control processes to determine if a change in strategy is warranted (Shenhav et al., 2016). Investigating whether control-related computation would similarly engage the cingulate’s intra-telecenphalic pathway, or whether it has a separate anatomical pathway for ultimately releasing the pyramidal tract from inhibitory influence, is an interesting avenue for future studies.

Finally, we note that our observation of a marked modulation of ACC ensemble representation associated with a specific action that the animal commits to, depending on whether that action is part of the ongoing strategy or is being re-evaluated as an alternative, may hint at what is encoded by the PT(CN) pathway. If, in an environment with a richer strategy space, this modulation further scales with the relative prevalence of a specific strategy, then this form of a representation would be empirically equivalent to a sampling from a probabilistic distribution over the strategy space (Fiser et al., 2010). Since our results argue that ACC’s PT(SNr) actively guides exploration of those strategies, demonstration of activity scaling with their local prevalence would suggest that strategies are sampled in accordance with this distribution; such a finding would lend credence to the idea that cingulate pyramidal tract pathway encodes a probabilistic internal model over strategy space.

## Acknowledgements

We thank Kim Ritola and the rest of Janelia viral core with help with high-titer virus production, Susan Michael for help with histology, and Maria Rysakova for help with tracing experiments. We thank Shaul Druckmann and Vivek Jayaraman for advice and comments on the manuscript. K. Vicari created the task illustration.

## Author Contributions

D.G.R.T., E.K. and A.Y.K. designed the behavioral training protocol, trained animals to perform the task and performed optogenetics experiments. M.K. built custom components and wrote software for the electrophysiology setup. M.K., E.K. and M.P performed the electrophysiology experiments. A.L. built custom components and wrote software for wireless optogenetics experiments. R.B. generated viral tools used in this study. M.M., D.G.R.T, E.K. and A.K. analyzed data. D.G.R.T, E.K. and A.K. wrote the manuscript.

## Data and materials availability

Requests for data and materials should be addressed to A.Y.K.

## Supplementary Figure Legends

**Figure S1, related to Figure 1.**
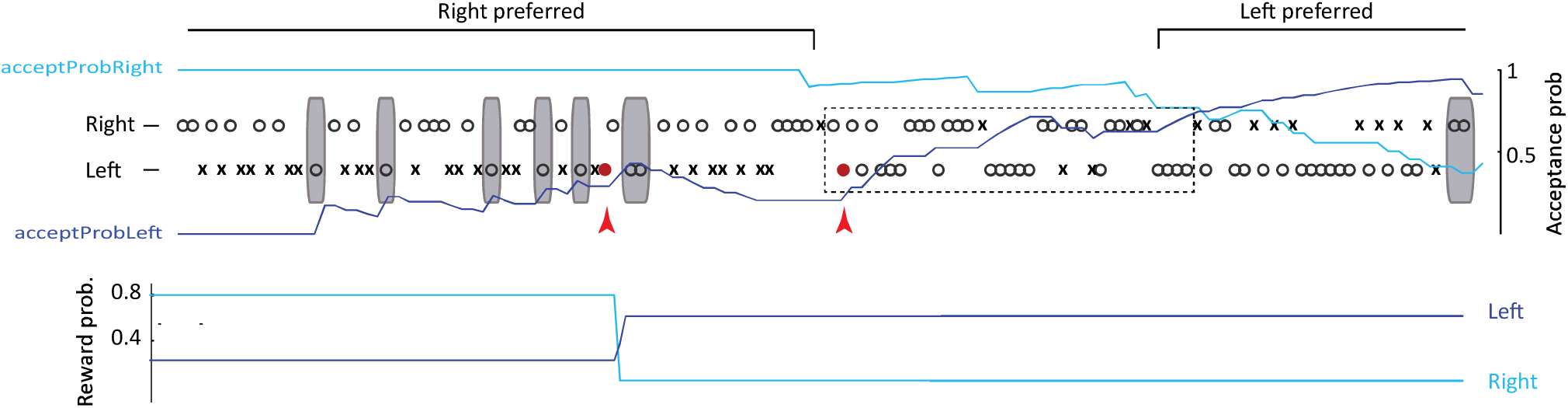
Sample behavioral trace from Figure 1B, with error trials included. **Top panel:** Sample behavioral trace around a block transition. Red errors indicate the two error trials (when the animal responded at the wrong lever) omitted for clarity from Figure 1B.

**Figure S2, related to Figure 1.**
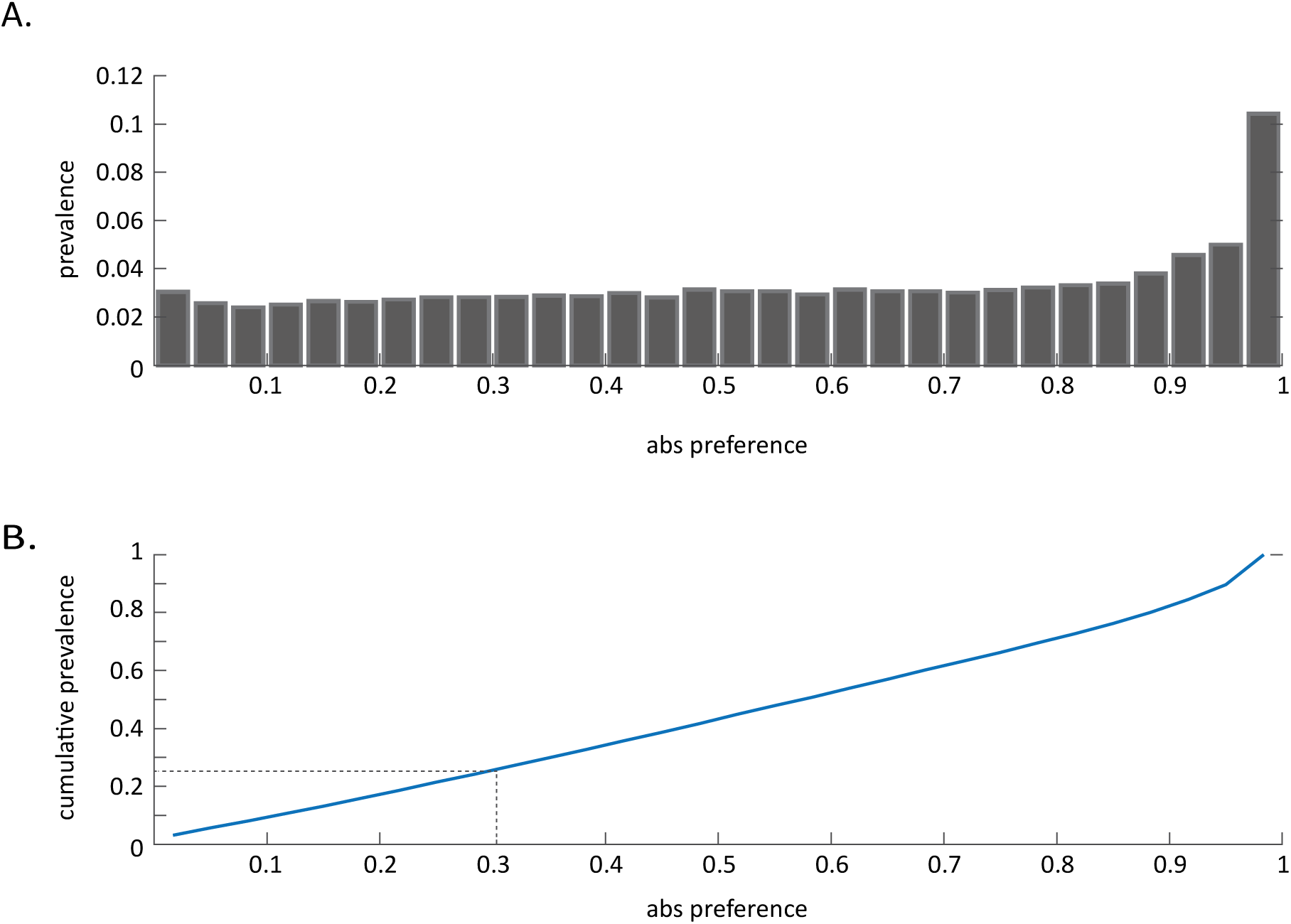
Periods of marked side preference dominate behavioral sessions. Prevalence **(A)** and cumulative prevalence **(B)** for different absolute values of behavioral preference (difference between local acceptance probability of the two choice options).Dashed line in **(B)** - threshold preference for inclusion into the analyses.

**Figure S3, related to Figure 1.**
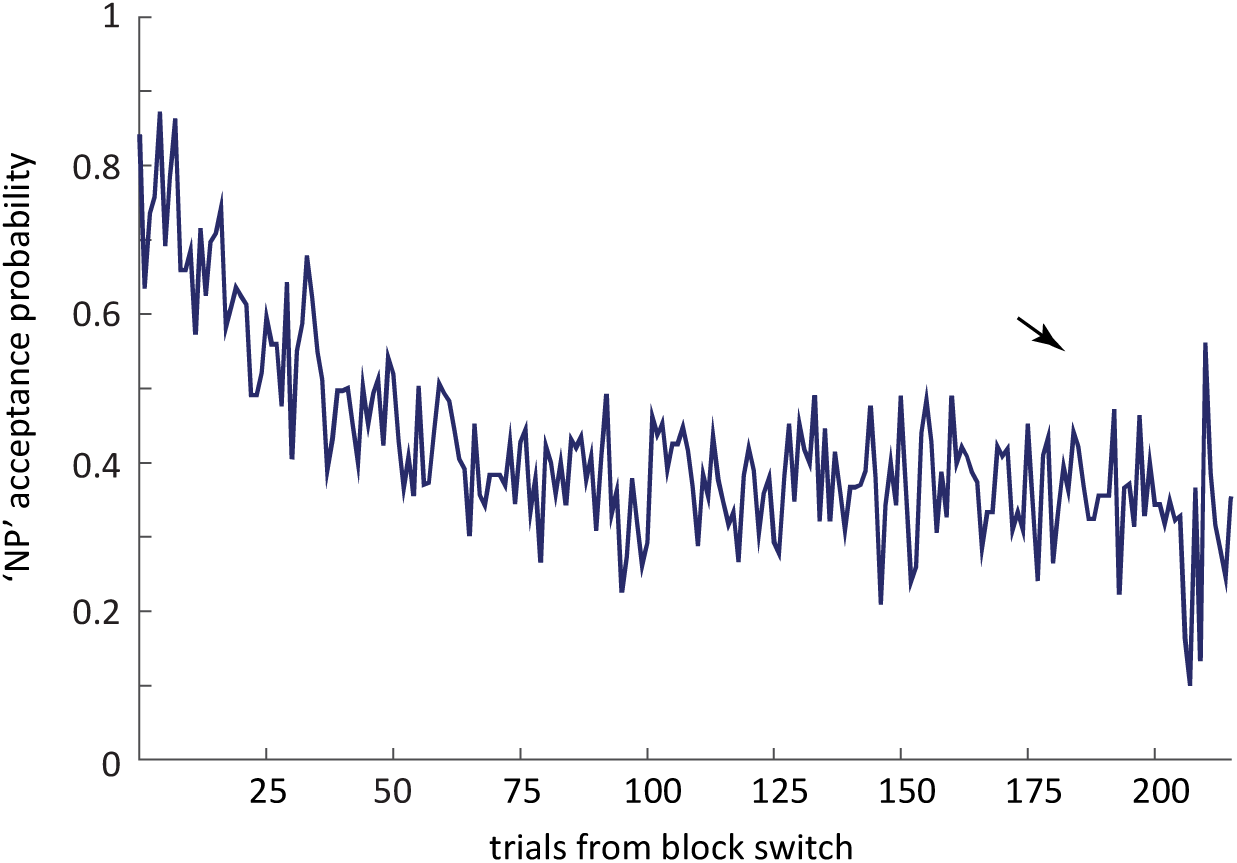
Sporadic decisions to accept the ‘non-preferred’ choice option persist late into blocks of stable reward probabilities. Local acceptance probability of choice option associated with smaller reward probability following block transition. Note sustained non-zero acceptance probability late into the block (arrow).

**Figure S4, related to Figure 1.**
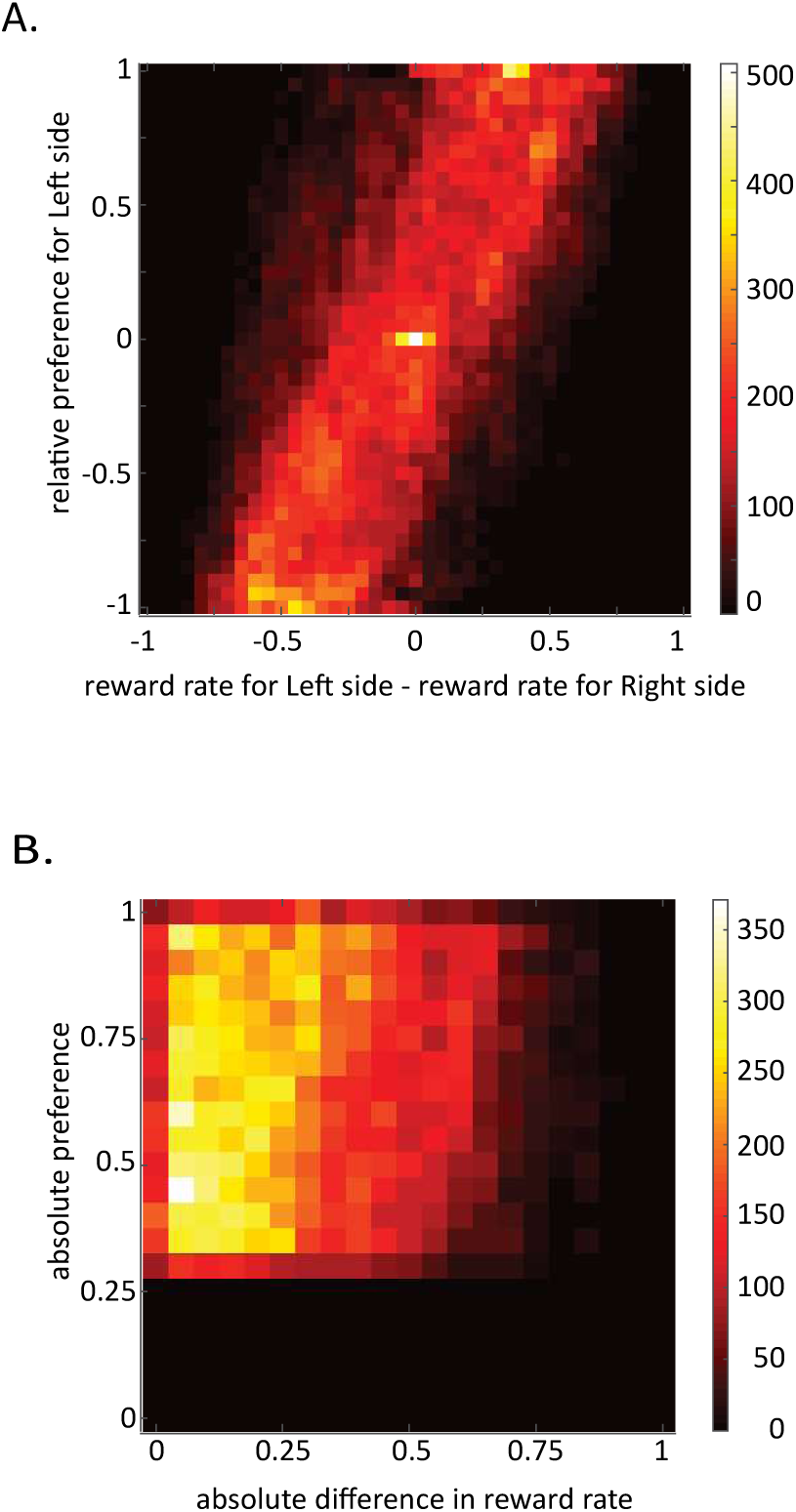
Animals sample the ‘non-preferred’ option irrespective of the relative reward rate. **A**, Relative preference for Left side as a function of the difference in reward rate between Left and Right sides. Note the extensive spread along the y axis even when the reward probabilities are equal. **B**, Behavioral preference as a function of the difference in reward rate for epochs under study (i.e when preference exceeds 0.3). Note little relationship between the relative rate of reward and the extent to which the ‘non-preferred’ side is sampled (acceptance probability of non-preferred choice option ∼ 1- relative side preference).

**Figure S5, related to Figure 1.**
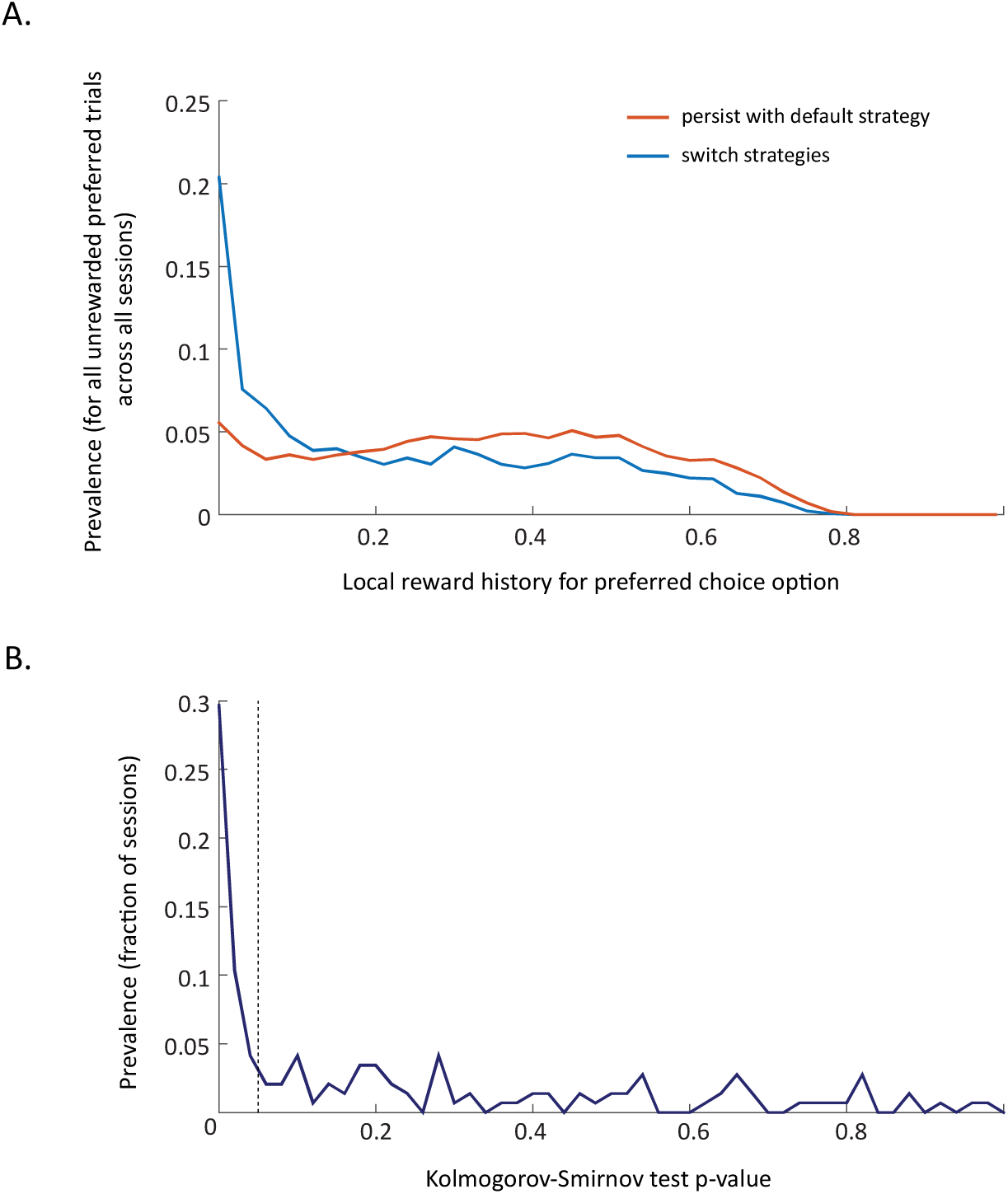
Reward history before decisions to persist (reject the subsequent ‘non-preferred’ choice option) or switch (accept the subsequent ‘non-preferred’ choice option) are often similar. **A**, Prevalence of local reward history values associated with the decisions to persist or switch in face of a transient downturn. All trials from all sessions are included. **B**, Prevalence of specific p-values for the Kolmogorov-Smirnov test for the difference between distributions as in A, when comparison is carried out within individual sessions.

**Figure S6, related to Figure 1.**
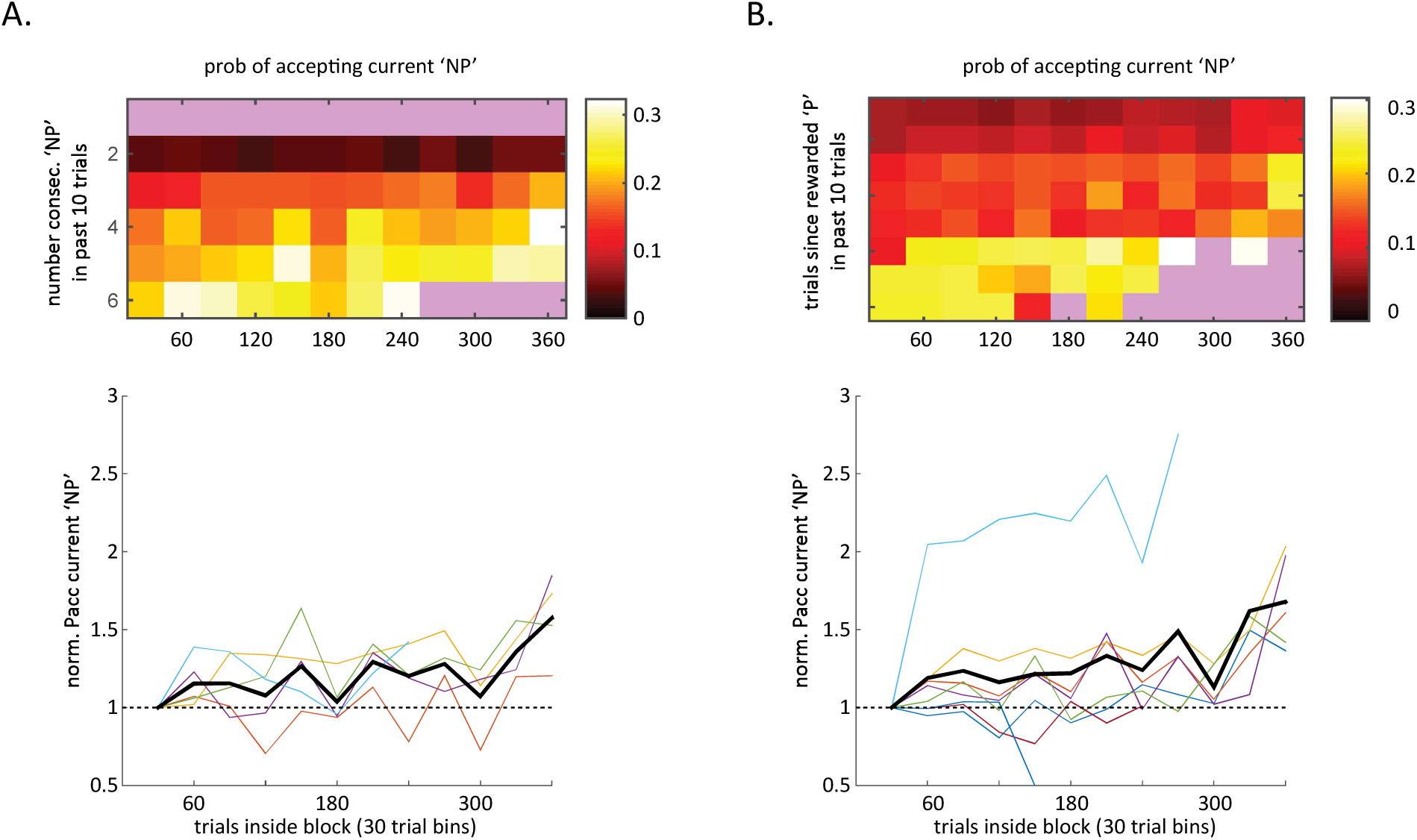
Decisions to accept the ‘non-preferred’ option reflect knowledge of block statistics. **A, Upper panel:** Heat plot of the prevalence of accept decisions for the ‘non-preferred’ choice option on offer as a function of the number of trials inside a block of stable probabilities (x axis, values estimated for 30 trial bins) and of the number of consecutive encounters the ‘non-preferred’ choice option within the past 10 trials (y axis). Black/white color denote no/frequent acceptances of the ‘non-preferred’ option respectively; purple color denotes values without sufficient data. **Lower panel:** The same data normalized to the initial prevalence of the acceptance of the ‘non-preferred’ option. Each colored line represents the normalized prevalence of accept decisions for the ‘non-preferred’ choice option on offer given different parameterizations of the consecutive encounters with it in the previous 10 trials. Black line represents the mean of these different parameterizations. **B**, Same as **A**, just for different parametrizations of the number of trials since the last reward was collected for the pursuit of the ‘preferred’ choice option.

**Figure S7, related to Figures 1,4.**
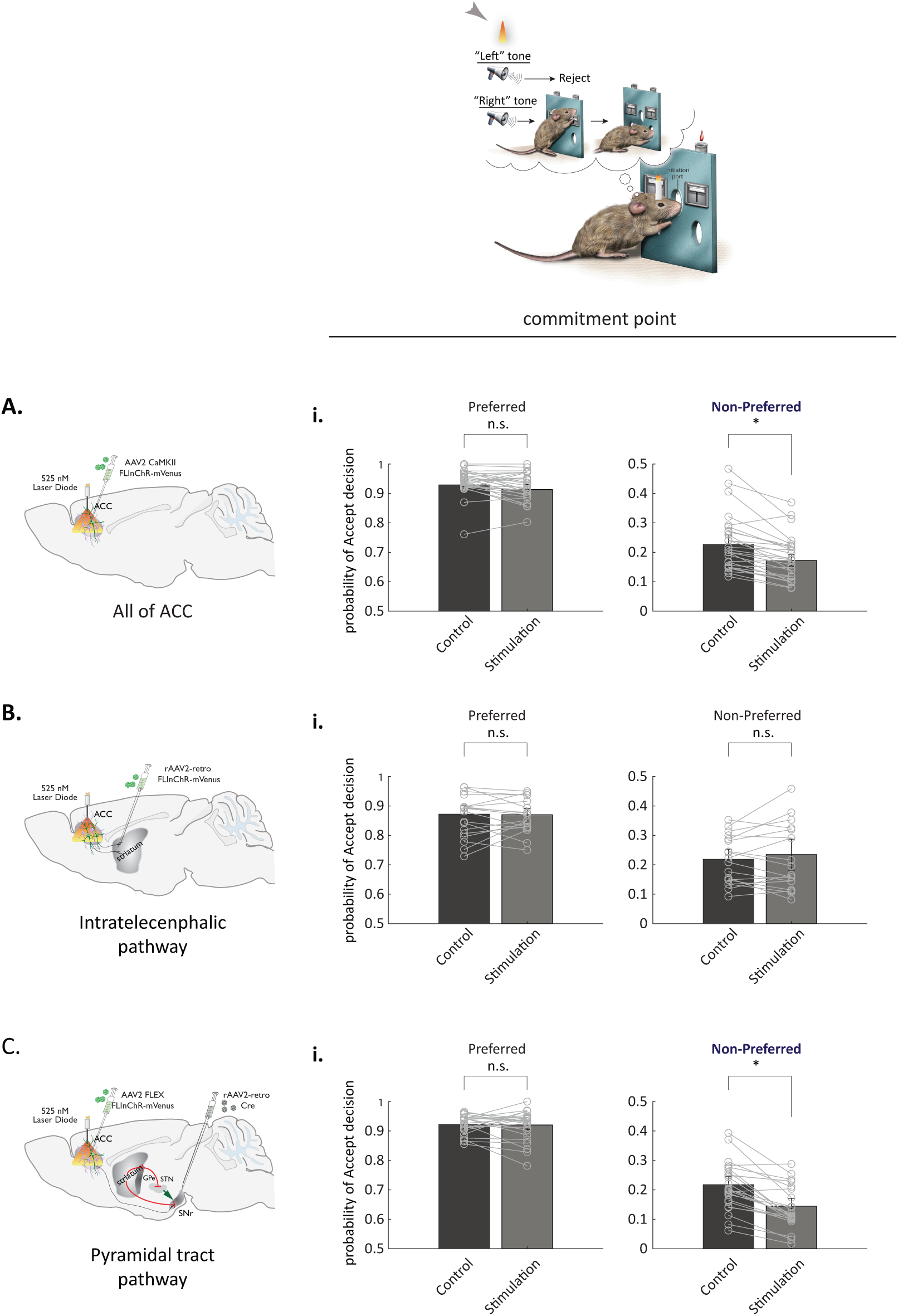
Perturbation of all of ACC, or of its PT-SN pathway, at the commitment point leads to decreased pursuit of the ‘non-preferred’ choice option. Absolute probability of accept decision for “preferred” **(i)** and “non-preferred” **(ii)** choice option in control condition and during transient Perturbation of all of ACC **(A)**, its IT-CS pathway **(B)** or its PT-SNr pathway **(C)** at the commitment point. Points represent individual sessions. Bars represent averages across within-animal means. Error bars represent standard error of the mean. n.s. non-significant, * p< 0.05, sign rank test with correction for multiple comparisons.

**Figure S8, related to Figures 1, 4.**
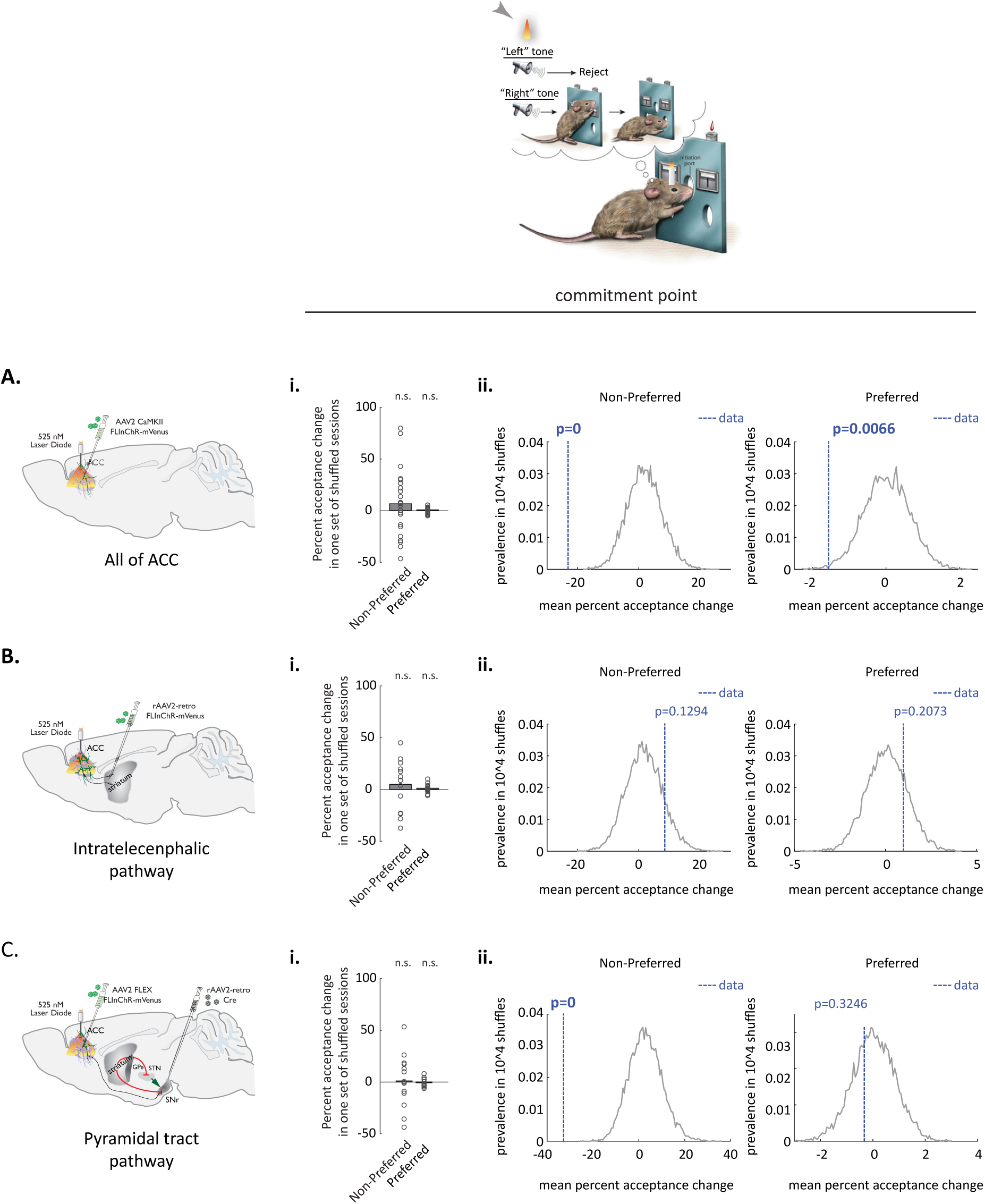
Observed changes in acceptance of the ‘non-preferred’ choice option following perturbation of ACC, or its PT-SN pathway, at the commitment point fall outside of the distribution of values obtained by chance. **(i)** Percent acceptance changes observed in a single set of shuffles, with ‘light-on’ vs ‘light-off’ labels randomly scrambled within each session in “all of ACC” **(A)**, “IT-CS” **(B)** and “PT-SN” **(C)** groups. Bars-across-session mean percent acceptance change for one set of shuffled sessions. **(ii)** Distribution of mean percent acceptance changes for the ‘non-preferred’ (left panel) and ‘preferred’ (right panel) choice options in 10,000 shuffles (the null distribution). Dashed blue lines-experimentally observed mean percent acceptance change. Note that large (25-40%) experimentally observed changes in acceptance of the ‘non-preferred’ choice option following perturbation of all of ACC (Aii, left panel), or of its PT-SN pathway (Cii, left panel) fall outside of the null distribution. Note also that the observed small change (<2%) in acceptance of the ‘preferred’ choice option following perturbation of all of ACC similarly falls outside of the null distribution (Aii, right panel). However, the underlying effect was not consistent across all animals (Figure S8), and is absent following perturbation of the PT-SN pathway (Cii, right panel).

**Figure S9, related to Figures 1,4.**
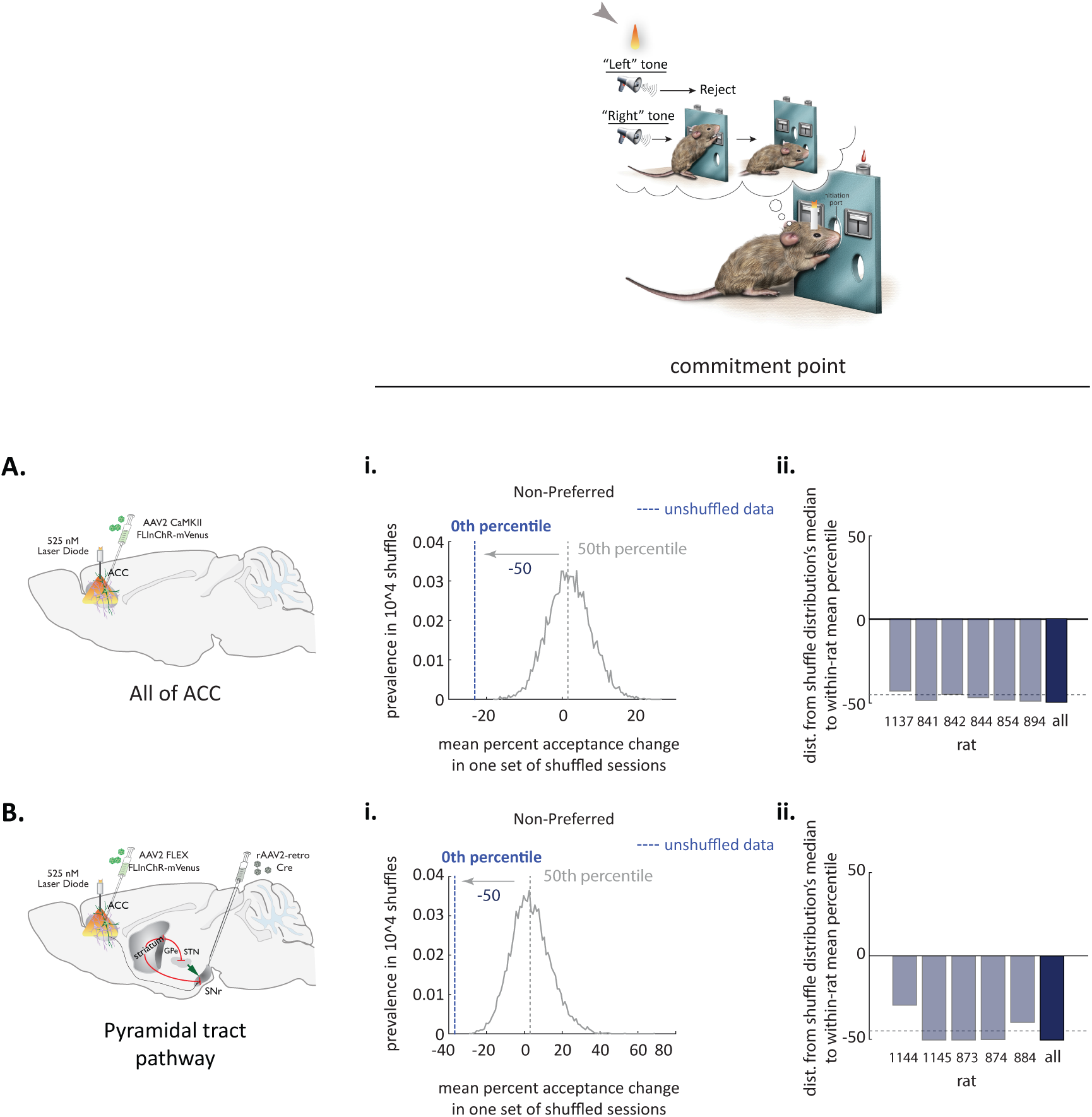
All animals contribute to pushing the experimental mean change in acceptance of the ‘non-preferred’ choice option following Perturbation of ACC, or its PT-SN pathway, at the commitment point outside of the null distribution. **(i panels)** Null distributions of mean percent acceptance changes for the ‘non-preferred’ choice option in 10,000 shuffles (replotted from Figure S7) for the Perturbation of all of ACC **(A)** or its PT-SN pathway **(B)**, with a schematic for the metric. Dashed blue lines-experimentally observed mean percent acceptance change (here, calculated across all animals). Dashed grey line-median (50th percentile) of the null distribution. Here, experimental mean fell to the left of the null distribution, so the metric (distance from the null distributrion’s median to across-rat mean percentile) is negative. **(ii panels)** Distances from the null distribution’s median to within-rat means.

**Figure S10, related to Figures 1,4.**
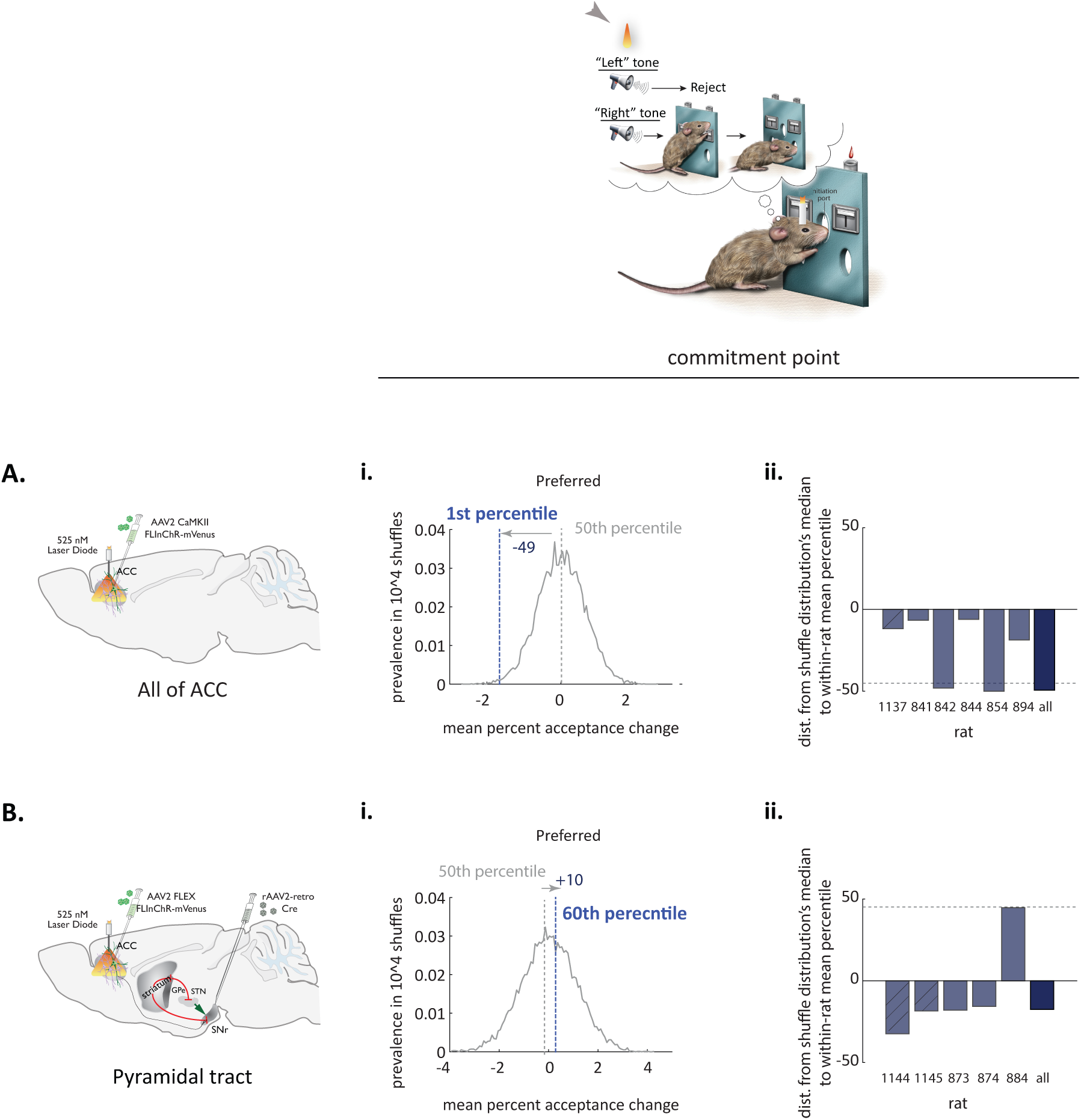
Effect of perturbing ACC (or its PT-SN pathway) on acceptance decisions for ‘preferred’ choice option is small and not consistent across animals. **(i panels)** null distributions of mean percent acceptance changes for the ‘preferred’ choice option in 10,000 shuffles (replotted from Figure S7) for the Perturbation of all of ACC **(A)** or its PT-SN pathway **(B)**, with a schematic for the metric. Dashed blue lines-experimentally observed mean percent acceptance change (here, calculated across all animals). Dashed grey line-median (50th percentile) of the null distribution. Here, experimental mean fell to the left of the null distribution, so the metric (distance from the null distribution’s median to across-rat mean percentile) is negative. **(ii panels)** Distances from the null distribution’s median to within-rat means. Dashed bars represent animals with the dual viral targeting strategy.

**Figure S11, related to Figure 1.**
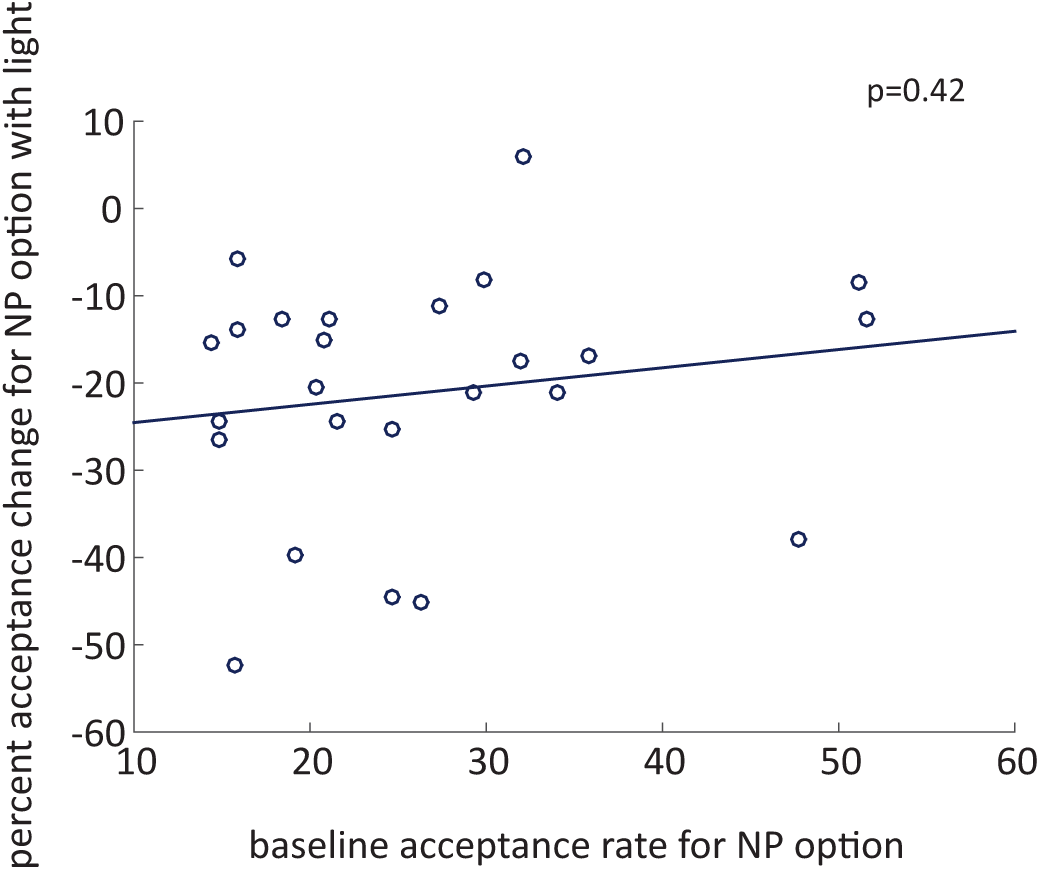
Perturbation effect size does not correlate with baseline incidence of accept decisions for the ‘non-preferred’ choice option. Illuminaon-dependent fractional change in the incidence of ‘accept’ decisions for the non-preferred (NP) option as a function of baseline rate of acceptance.

**Figure S12, related to Figures 3,4.**
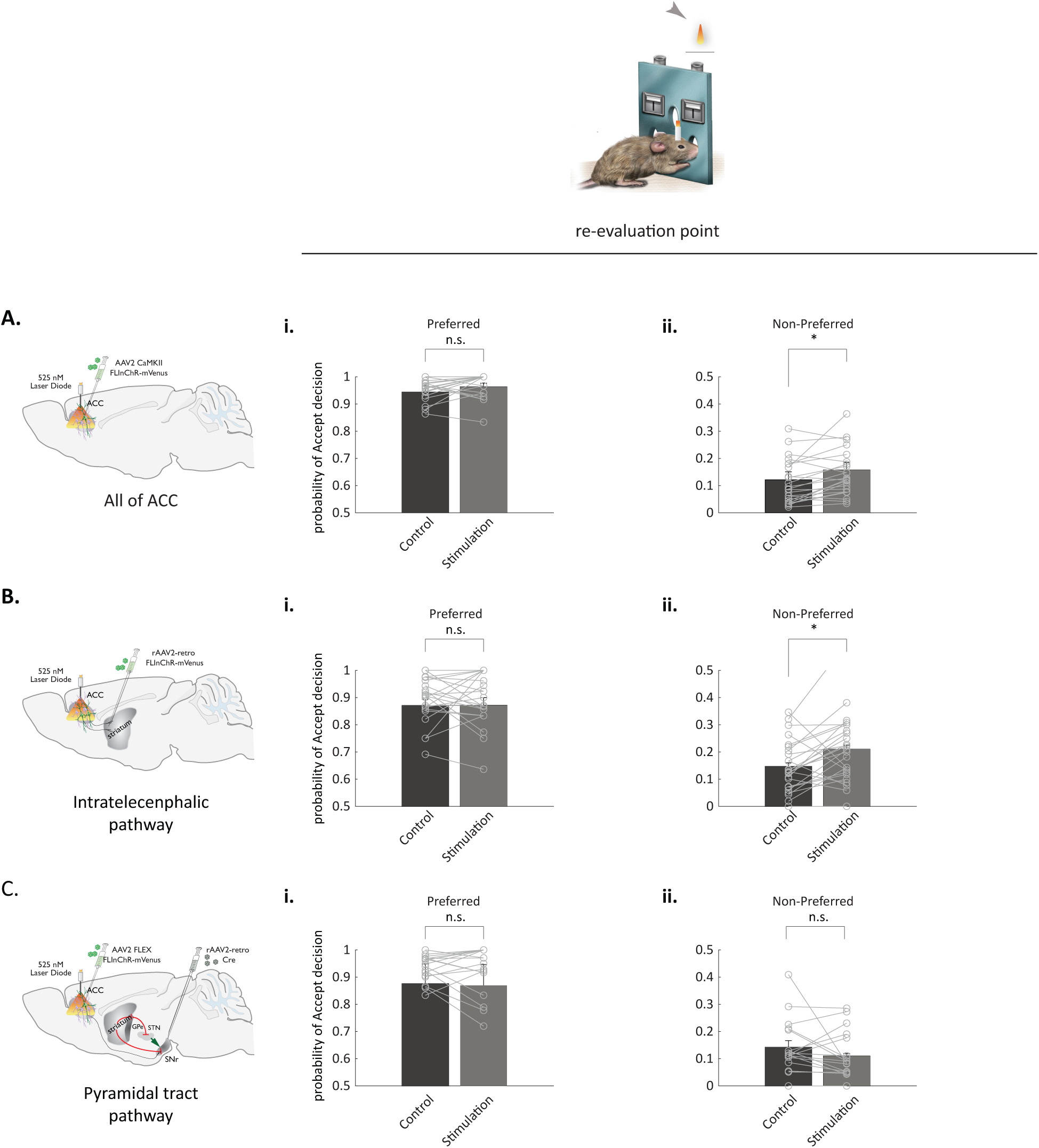
Perturbation of all of ACC, or of its IT-CS pathway, at the re-evaluation point leads to increased pursuit of the ‘non-preferred’ choice option. Absolute probability of accept decision for “preferred” **(i)** and “non-preferred” **(ii)** choice option in control condition and during transient Perturbation of all of ACC **(A)**, its IT-CS pathway **(B)** or its PT-SNr pathway **(C)** at the commitment point. Points represent individual sessions. Bars represent averages across within-animal means. Error bars represent standard error of the mean. n.s. non-significant, * p< 0.05, sign rank test with correction for multiple comparisons.

**Figure S13, related to Figures 3,4.**
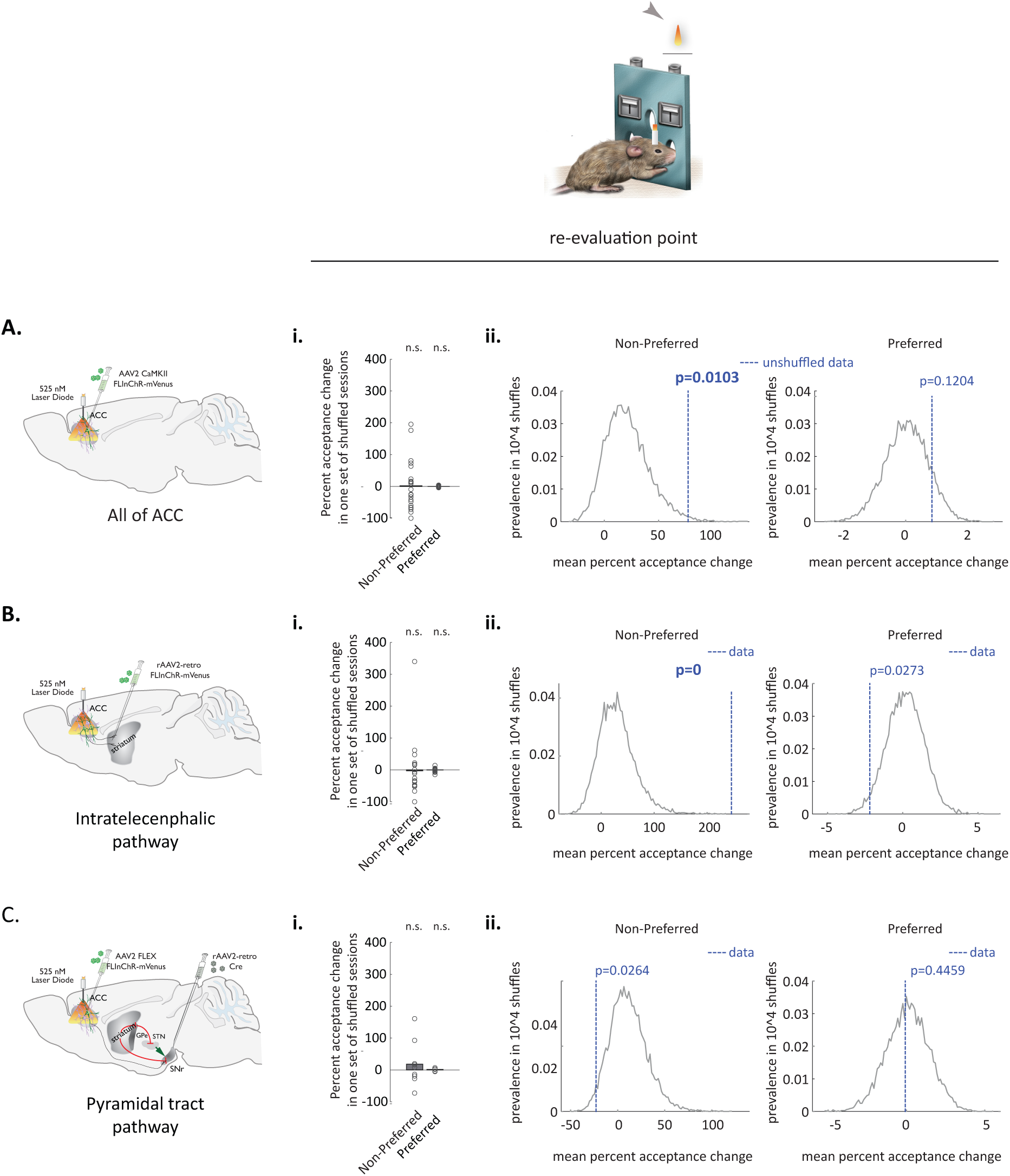
Observed changes in acceptance of the ‘non-preferred’ choice option following Perturbation of ACC, or its IT-CS pathway, at the re-evaluation point fall outside of the distribution of values obtained by chance. **(i)** Percent acceptance changes observed in a single set of shuffles, with ‘light-on’ vs ‘light-off’ labels randomly scrambled within each session in “all of ACC” **(A)**, “IT-CS” **(B)** and “PT-SN” **(C)** groups. Bars-across-session mean percent acceptance change for one set of shuffled sessions. **(ii)** Distribution of mean percent acceptance changes for the ‘non-preferred’ (left panel) and ‘preferred’ (right panel) choice options in 10,000 shuffles (the null distribution). Dashed blue lines-experimentally observed mean percent acceptance change. Note that large (70-90%) experimentally observed changes in acceptance of the ‘non-preferred’ choice option following Perturbation of all of ACC (Aii, left panel), or of its IT-CS pathway (Bii, left panel) fall outside of the null distribution. Note that for significance, the p value would need to be less than 0.025 given the two separate tests (for preferred and non-preferred trials) done for each dataset.

**Figure S14, related to Figures 1,4.**
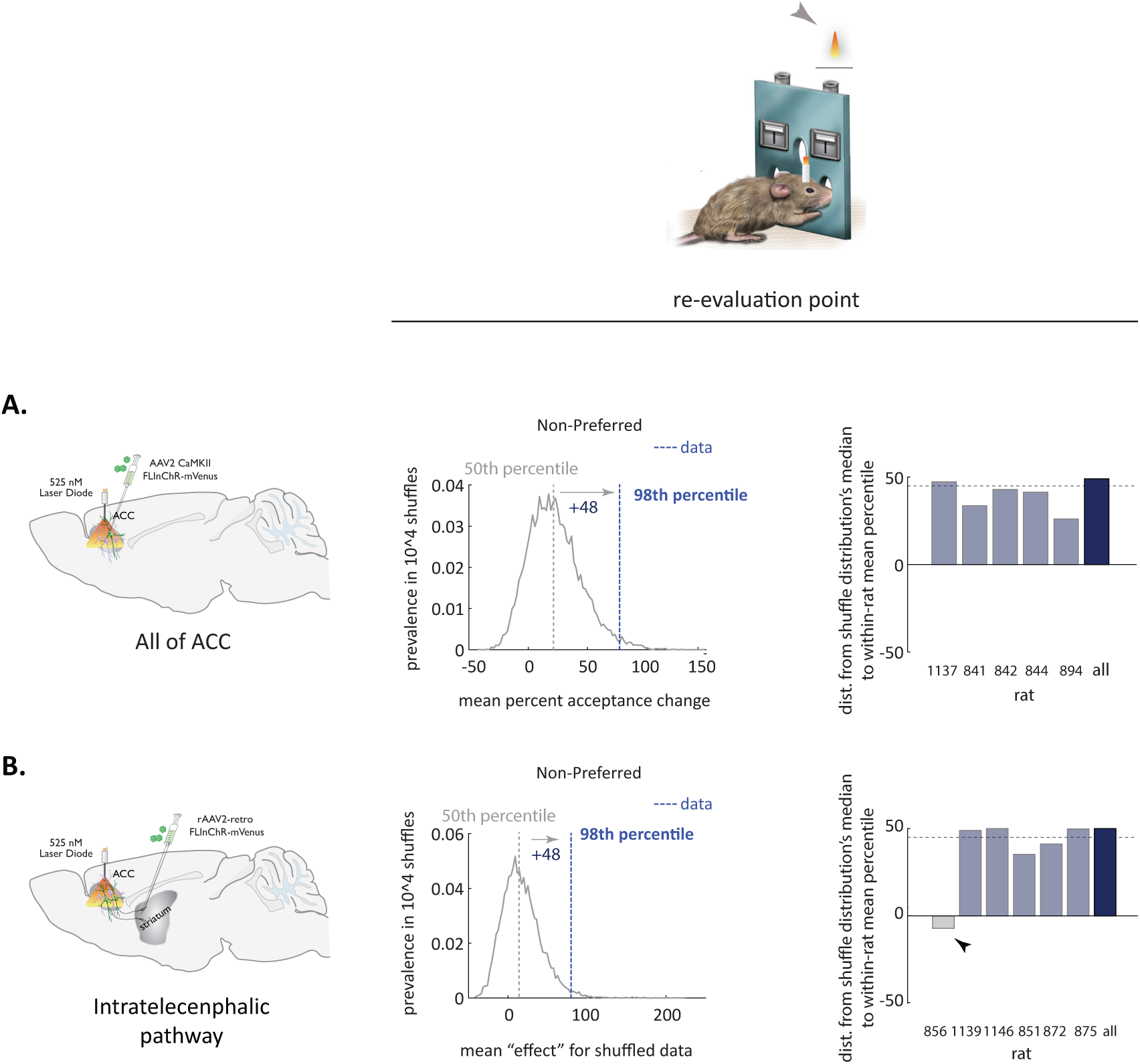
All properly targeted animals contribute to pushing the experimental mean change in acceptance of the ‘non-preferred’ choice option following Perturbation of ACC, or its IT-CS pathway, at the re-evaluation point outside of the null distribution. **(i panels)** Null distributions of mean percent acceptance changes for the ‘non-preferred’ choice option in 10,000 shuffles (replotted from Figure S11) for the Perturbation of all of ACC **(A)** or its IT-CS pathway **(B)**, with a schematic for the metric. Dashed blue lines-experimentally observed mean percent acceptance change (here, calculated across all animals). Dashed grey line-median (50th percentile) of the null distribution. Here, the experimental mean fell to the right of the null distribution, so the metric (distance from the null distribution’s median to across-rat mean percentile) is positive. **(ii panels)** Distances from the null distribution’s median to within-rat means. Arrow in **Bii** points to the rat, for which targeting of the viral delivery failed (see Figure S13). Dashed bars represent animals added in revision.

**Figure S15, related to Figures 4, S12.**
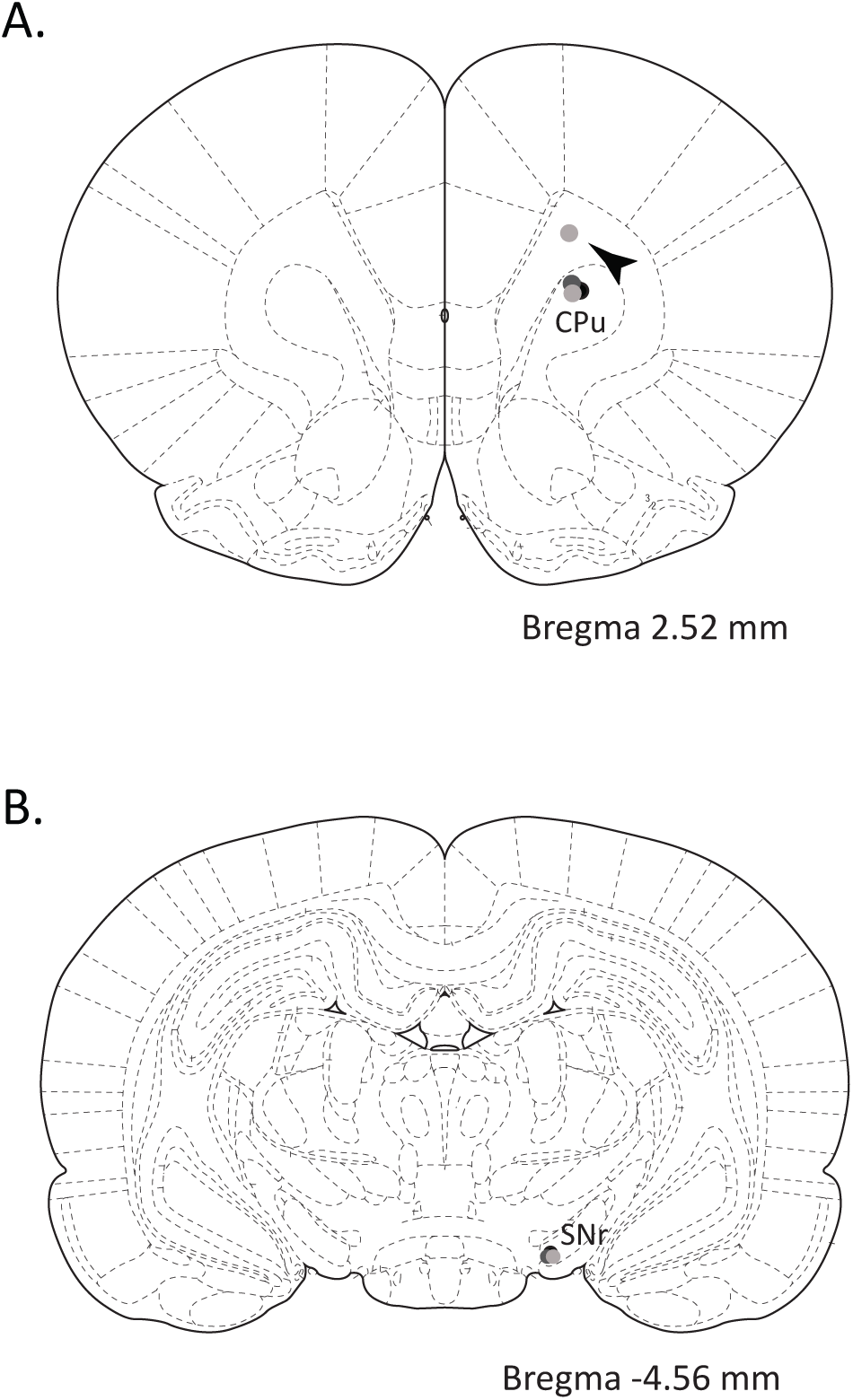
Locations of viral injections into the axonal fields. Locations of injections in individual animals for the groups targeting the IT(CS) pathway **(A)** or the PT(CN) pathway **(B)**. Arrow in **(A)** points to one rat, for which the targeting failed. See Figure S12.

**Figure S16, related to Figure 4.**
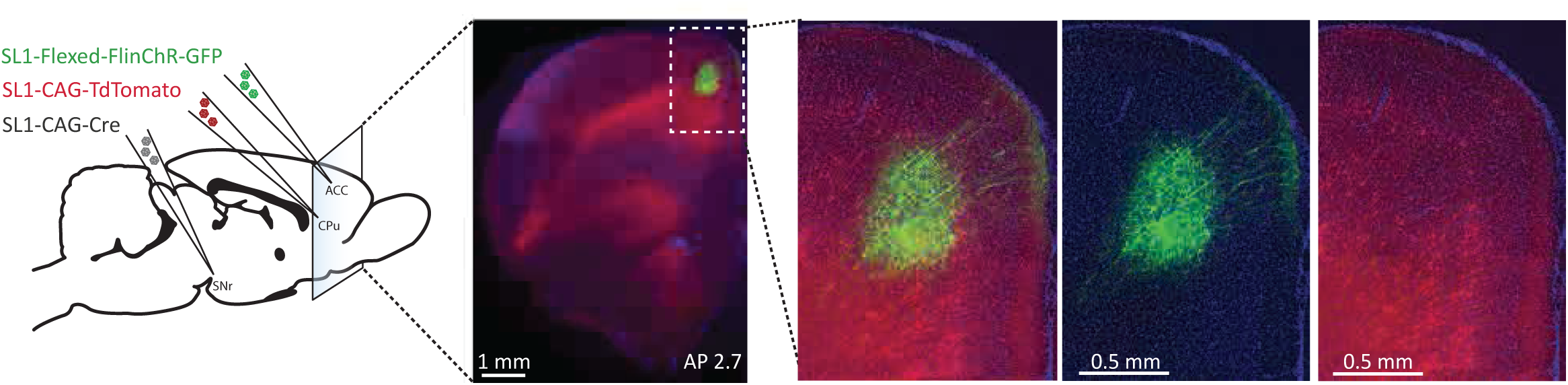
PT(CN) pathway is not accessible through striatal injections. Double labeling of the PT(CN) pathway in green and of the IT(CS) pathway in red.

## Experimental Procedures

### Subjects

All experiments were done in male Long Evans rats (400-500g). Animals were kept at 85% of their initial body weight before food restriction and maintained on a 12hr light/12hr dark schedule. Experiments were conducted according to National Institutes of Health guidelines for animal research and were approved by the Institutional Animal Care and Use Committee at HHMI’s Janelia Farm Research Campus.

18 animals were implanted for optogenetic experiments (6 each targeting all of ACC excitatory neurons, the cortico-striatal population, and the cortico-nigral population). One of the cortico-nigral animals had to be sacrificed prior to collection of any data due to poor recovery from the surgery. All remaining animals were used for perturbation experiments. With one exception, the same set of animals in each of the three experimental group (“all of ACC”, “IT-CS” and “PT-CN”) was used for perturbation experiments at the point of commitment and the time of strategy re-evaluation. One animal in “all of ACC” group developed an infection around the implant and had to be euthanized early on in the multi-week experiment thus contributing data only to the set of perturbations at the initiation.

1 animal was implanted with a tetdrode drive for collection of neural activity during the task. Data from this animal was combined with data from 3 animals used in a previous study (Karlsson et al., 2012).

### Behavioral apparatus and task

All behavior was confined to a box with 23 cm high plastic walls and stainless-steel floors (Island Motion Corp). The floor of the box was 25cm by 34 cm, and the levers and nose ports were all arranged on one of the 25 cm walls. All lights, nose ports, levers, and reward deliveries were controlled and monitored with a custom-programmed microcontroller, which in turn communicated via USB to a PC running a Matlab-based control program. Nose port entries were detected with an infrared beam-break detector (IR LED and photodiode pair). The central initiation port contained one white LED that indicated the option to initiate a new trial. The left and right levers were pneumatically extracted from the wall upon each trial initiation and were retracted after one of the two levers was pressed. Simultaneously upon initiation, one of two sounds was presented across two speakers (located on the two 34cm walls) with equal volume, and they were frequency modulated (1% modulation at 6.67 Hz) around a single base frequency. A 6.5 kHz base frequency signified that the left lever was correct, and a 14 kHz base frequency signified that the right lever was correct. Either tone had an equal chance of being presented upon initiation. Tone presentation was terminated after 2 second or upon the first lever press, whatever came first. All behavior was video recorded at 30 frames/sec using an infrared-sensitive camera. Except after incorrect lever presses, the only visible light source inside the box was from the LED in the initiation port. During error trials, a white LED array in the ceiling of the box was lit, and no trials could be initiated during a timeout period. Liquid rewards (0.1 ml drops of 10% sucrose mixed with black cherry Kool-Aid) were delivered from the reward ports 0.5 seconds after port entry with a motorized syringe pump (Harvard Apparatus PHD 2000).

### Behavioral training

Food-restricted animals were trained to perform the task with minimal ‘shaping’—from the first moment of training, animals were exposed to the full task with four exceptions: 1) animals only needed to press the correct lever once before reward became available, 2) reward probabilities were kept at or above 0.5 for both sides, 3) the timeout period for pressing the wrong lever was kept short (0.5 seconds) and 4) the time that was allowed to pass between initiating the trial and pressing the lever, as well as between pressing the lever and collecting the reward was 300 sec. At first, animals performed the correct sequence of actions rather infrequently and by chance. However, after approximately 1-2 weeks of training, most animals learned the entire task structure, including the presence of the option to reject trials and of unsignaled changes in reward contingencies. After each animal successfully completed two trial blocks in one session, the number of required lever presses was gradually increased to 5, the timeout period was increased to 30 seconds, and reward probabilities became randomly drawn from a set spanning low and high values (0.2, 0.3, 0.4, 0.5, 0.6, 0.7 and 0.8) at block transitions. Reward probabilities associated with the two sides were selected independently. This design feature made it difficult for the rat to infer the identity of the new and more profitable option simply from the change in reward probability of the preferred option, thus prompting exploratory bouts.

Animals were considered proficient on the task when they preferentially rejected the less profitable side and dynamically updated this rejection policy when reward probabilities changed. 90% of animals became proficient within one month of training. To prevent animals from predicting when reward probability changes would occur, the number of trials within each block was drawn randomly from a Gaussian distribution (mean 250 with standard deviation of 25). Other than the changed reward probabilities, no cues were presented to indicate block transitions.

### In vivo virus injections

For localized *in vivo* viral delivery, rats were anaesthetized with isoflurane (∼2% by volume in O_2_; SurgiVet, Smiths Medical) and a small hole was drilled in the skull above the requisite injection site (see table below). For some injection sites, several injections were made at different depths (see table below). For viral injections, ∼ 250-500 nl of virus-containing solution was slowly injected at each depth into the tissue. Injections were done with a pulled glass pipette (broken and beveled to 25–30 μm (outside diameter); Drummond Scientific, Wiretrol II Capillary Microdispenser) back-filled with mineral oil. A fitted plunger was inserted into the pipette and advanced to displace the contents using a hydraulic manipulator (Narashige, MO-10). Retraction of the plunger was used to load the pipette with virus. The injection pipette was positioned with a Sutter MP-285 manipulator.

The following coordinates were used in this study:

| Injection target | Injection coordinates, mm* |
| --- | --- |
| ACC (Cg1 (Vogt and Paxinos, 2014)) | A/P: 2.50; M/L: 0.60; D/V: -2.42, and -2.1 |
| dorsomedial striatum (DMS) | A/P 2.16; M/L 1.7 ; D/V -4.3 |
| SNr | A/P -4.75; M/L ±2; D/V -8.6, -8.55 |

\* All A/P coordinates are given with respect to Bregma. All D/V coordinates are with respect to the skull.

The following viral constructs were used:

| Perturbation target | Viral constructs, injection location | Fiber location |
| --- | --- | --- |
| All of ACC | AAV2/1 pCAMKII- FLInChR-mVenus into ACC | ACC |
| Cortico-striatal pathway | AAV2-retro pCAG- FLInChR-mVenus into DMS (4 animals)<br>AAV2-retro pSyn-Cre into DMS,<br>AAV2/1 FLEX-CAG-FLInChR-mVenus into ACC (2 animals) | ACC |
| Corticonigral | AAV2-retro pSyn-Cre into SNr,<br>AAV2/1 FLEX-CAG-FLInChR-mVenus into ACC | ACC |

### Wireless optogenetics system

The “Cerebro” optogenetics system consists of an implantable programmable device that is triggered remotely by infrared light. The implanted component comprises two laser diodes (520nm; Cat No: PL520B1; World Star Tech) coupled to a pair of 200um core, 0.5NA hard polymer optic fibers (FP200URT; ThorLabs) cut to protrude from the acrylic base of the implant by approximately 2.5mm. A photoresistor is used as feedback to maintain a constant light output. Full manufacturing and parts information are provided online (https://karpova-lab.github.io/cerebro/).

To improve light scattering from the optical fibers, two methods were deployed. For some animals, we bonded 106-125um diameter silver spheres (Cat. No. M-60-0.17, CoSpheric) to the tip of these fiber optics to scatter emitted laser light more uniformly throughout neuropil. For others, we sharpened the fibers using chemical etching approach(Hanks et al., 2015). Specifically, the cable jacket, strengthening fibers, and outer plastic coating (typically white or orange) were fully removed, leaving 1 cm of fiber optic cable and inner plastic coating intact. Then 2 mm of the fiber tip (with final layer of plastic coating still attached) was submerged in 48% hydrofluoric acid topped with mineral oil for 85 min, followed by water for 5 min (submerging 5 mm), and acetone for 2 min (to soften the plastic). The plastic coating was then gently cut with a razor and pulled off with tweezers to reveal a 1 mm sharp-etched fiber tip.

In a subset of animals, the module was implanted during viral injection surgery. In others, the implantation was done as a separate surgery, 3-4 weeks after viral delivery.

### Perturbation experiments

To avoid adaptation to perturbation, experiments were run at most every other day, and at times were interspersed with behavioral sessions where no light was delivered.

For perturbation at the time of trial initiation (commitment point), light delivery was done on 25% of all trials. The laser diode was turned on immediately upon central nose port entry, and the delivery of the auditory tone was delayed by 50msec. If the rat rejected the trial, the laser light waveform was immediately terminated. If the rat accepted the trial, the light waveform was permitted to progress to conclusion (2 sec), and then ramped down.

For perturbation during strategy re-evaluation, light delivery was done on 15-35% of all completed trials; a 2 sec waveform was triggered upon reward port entry. Since animals typically licked at the reward port even on unrewarded trials, the 2 second perturbation period did not typically bleed into the following trial. On rare occasion when animals did proceed to initiate the next trial in less than 2 sec, the light waveform was immediately terminated by central port entry.

The two experimental protocols were carried out in different experimental sessions.

### Electrophysiological recordings

Data from 3 out of 4 animals were reported previously(Karlsson et al., 2012) and the associated methods were described in detail in that manuscript. The methods for the fourth animals followed a same paradigm except for a different angle of approach and a new recording system. Briefly, a microdrive array containing 20 independently movable tetrodes(O’Keefe and Recce, 1993) was chronically implanted on the head of the animal. Each tetrode was constructed by twisting and fusing together four insulated 13 μm wires (stablohm 800A, California Fine Wire). Each tetrode tip was gold-plated to reduce impedance to 200-300 kΩ at 1 kHz. Within the implant, the tetrodes converged to a circular bundle (1.9 mm diameter), angled either at 20° or at 0° with respect to vertical (pointing towards midline after implantation). Over the two weeks following surgery, the tetrodes were slowly lowered to a depth of 2 mm along the 20° trajectory (for the first three animals), or vertically (for the fourth animal), moving approximately 160 μm/day on average. During this time, animals were re-acclimated to performing the task with the drive. When performance on the task was regained to pre-surgery levels (in terms of motivation and dynamic strategy arbitration behavior), recording sessions began. After each recording session, any tetrodes that did not appear to have any isolatable units were moved down 80 μm.

Each recording session spanned 1.5 to 4 hours, depending on the animal’s motivation to perform. Animals were not forced to perform the task and sometimes took breaks (generally around 5 minutes, but sometimes up to 30 minutes). Data from the first three animals were collected with an Nspike data acquisition system (L. Frank, UC San Francisco, and J. MacArthur, Harvard Instrumentation Design Laboratory). Data from the fourth animal was collected using the wireless headstage and datalogger (M. Karlsson, HHMI and SpikeGadgets). An infrared diode array with a large and a small group of diodes was attached to the animal’s preamplifier array and the animal’s position in the environment was reconstructed using a semi-automated analysis of digital video of the experiment with custom-written software.

Spike data were sampled at 30 kHz, digitally filtered between 600 Hz and 6 KHz (2 pole Bessel for high and low pass) and threshold crossing events were saved to disk. Continuous local field potential data from all tetrodes was sampled at 1.5 KHz, digitally filtered between 0.5 and 400 Hz and saved to disk.

Individual units on each tetrode were identified by manually classifying spikes using polygons in two-dimensional views of waveform parameters (Matclust, M.K.). For each channel of a tetrode, peak waveform amplitude and the waveform’s projection onto the first two principal components were used for clustering. Autocorrelation analyses were done to exclude units with non-physiological single-unit spike trains. Only units where the entire cluster was visible throughout the recording session were included.

Responses from a total of 381 units were analyzed in this manuscript.

### Data analysis

All data analysis was performed using custom-written routines in Matlab.

#### Identification of ‘preferred’ and ‘non-preferred’ trials

Iterated presentation of the two trial types (‘left’ and ‘right’) permitted us to make an independent calculation of the running acceptance probability for each behavioral option. pAcceptLeft and pAcceptRight were calculated with a sliding causal half-exponential (decay constant of 0.1), by assigning rejected trials a value of 0 and accepted trails the value of 1. Error trials not included in the calculation.

A trial was deemed ‘preferred’ (‘non-preferred’) if the acceptance probability for its cognate choice option was at least (at most) 0.5, and if the concurrent acceptance probability of the other choice option was smaller (greater) by at least 0.3. While all the results were robust to the choice of the latter threshold (the minimum difference in acceptance probability for the two sides), the difference of 0.3 was chosen to balance the desired focus on epochs with a markedly pronounced default strategy (i.e. side preference) and the need to have a sufficient sample size in the ‘non-preferred’ category.

Periods of preference within one session were pooled and analyzed together.

#### R.O.C. analysis of features predicting acceptance of non-preferred option

To determine whether, and how well, various behaviorally-relevant features (those present in the task and in the recent behavioral record) predicted the animals’ accept decisions for the ‘non-preferred’ choice option, we used a version of an ROC analysis that that uses the true negative and true positive rates to summarize how well a predictor dichotomizes a data set, summarized by the Youden’s Index, a R.O.C. analysis that uses the true negative and true positive rates to summarize how well a predictor dichotomizes a specific data set (Youden, 1950). Periods of marked side preference (see above) from all 160 behavioral sessions were stitched before the ROC analysis.

The behavioral features we examined fell into three main categories: the parameters associated with an individual trial leading up to the choice to be explained (at lags of 1-10 trials), running averages of different types of trials in recent past, and the number of trials since the last occurrence of a particular type of trial. Within the first category, we examined the following features: the type of choice option presented on that trial; whether the trial was accepted/rejected/an error trial; whether the trial was rewarded, whether the trial was both ‘preferred’ and rewarded, both ‘non-preferred’ and rewarded. Within the second category, we calculated **running averages of**: errors, presented choice options, acceptance probability for the ‘preferred’/’non-preferred’ choice option, and reward probability for ‘preferred’/’non-preferred’ choice option. In addition, we examined features that reflected **the number of trials** since the last reward, since the last reward for the ‘preferred’/’non-preferred’ choice option, since last acceptance, since last acceptance of the ‘preferred’ / ‘non-preferred’ choice option since last rejection, since last rejection of the ‘preferred’/ ‘non-preferred’ choice option, since the last presentation of the ‘preferred’/’non-preferred’ choice option presentation, since the beginning of the estimated start of a stable side preference, and since the beginning of a reward block. Finally, we looked at the experienced reward rates within each block, and the absolute difference in rejection of the ‘preferred’ and ‘non-preferred’ choice options. Each feature was examined within the space of all trials, and, when appropriate separately, within the subspace of trails of the relevant type.

Our final analysis (Figure 1C) focused on estimating the Youden’s Index for three features that performed significantly better than others at capturing the decisions to accept the ‘non-preferred’ choice option: number of trials (in the space of all trials) since the last rewarded ‘preferred’ trial, local probability of accepting the ‘non-preferred’ choice option (in the subspace of the ‘non-preferred’ trial type), and the number of consecutive encounters with the ‘non-preferred’ choice option (equivalent to the number of trials since the last ‘preferred’ choice option). For comparison, Youden’s Index was also calculated for one of the lesser-predictive features, local error rate.

#### Effect of environmental volatility on acceptance of the ‘non-preferred’ choice option

The capacity of local features to predict animals’ choices suggests that, in principle, a purely local strategy have been employed. To determine if, instead, higher-order task variables also influence animals’ acceptance of the ‘non-preferred’ choice option, we examined whether the prevalence of such decisions changed with the number of trials into a block of stable reward probabilities. To uncover the influence of such more global features in face of a strong contribution from the local ones, we evaluated how the prevalence of accepting the ‘non-preferred’ option changed as a joint function of the relevant global feature – time since the block start – and of individual local features identified above. Specifically, we plotted the prevalence of accepting the ‘non-preferred’ choice option as a function of the block depth (in 30-trial bins; normalized by the prevalence in bin 1) and of the different parameterizations of the local features (calculated for the 10-trial window). Similarly to the ROC analysis above, this analysis was performed on the entire behavioral dataset as a whole, with periods of marked behavioral preference stitched together across all sessions.

#### Analysis of optogenetic perturbation experiments

We calculated the ratio of acceptance probability between ‘light-on’ and ‘light-off’ conditions within each session for the trial type of interest (Figures 1,3 and 4). Statistics on the observed ratios was done across within-animal means. To guard against battery run-down in the optogenetic module, the upper limit on the number of trials used in the analysis was set at 1200 for the sessions with the perturbation during the commitment point, and to 1500 for sessions with perturbation during strategy re-evaluation point.

#### Shuffle analysis

To eliminate the possibility that the difference in the distribution of accept decisions for ‘light-on’ and ‘light-off’ trials occurred by chance, we repeated our analysis after randomly shuffling the ‘light-on’ and ‘light-off’ labels within individual sessions. If our observations arose by chance, such a manipulation would be expected to frequently produce values of mean fractional change in acceptance on the order of what we observed experimentally. Within each shuffle run, ‘light’ and ‘non-light’ labels would be independently shuffled for each individual experimental session for an individual condition (e.g. “all of ACC” @ commitment point). For perturbation sessions, during which the light stimulation occurred on initiation trials, any trial in the synthetic shuffle sessions could be designated as a light stimulation trial. For sessions, in which the light stimulation occurred during strategy re-evaluation, only trials that were not errors and were accepted could acquire the ‘light’ label to ensure that the synthetic shuffle sessions randomly distributed synthetic light stimulation on these correct, accepted trials. The change in accept decisions for ‘non-preferred’ and ‘preferred’ option as a result of this synthetic ‘light’ delivery was then computed across that run of generating synthetic session in the same manner as was done for the experimental data. This procedure was repeated 10, 000 times, and the “null distribution” of such synthetic mean percent changes was then constructed. The percentile value of experimentally observed mean percent change inside this ‘null’ shuffle distribution was calculated as the measure of significance of the result. When shuffle effects were evaluated for individual animals, the null distribution was constructed separately in each case, only using data for that animal.

#### Decoding analysis

For each session, we trained a linear classifier to predict, on the basis of ensemble activity, whether a particular trial was ‘preferred’ (part of the ‘default’ strategy) or non-preferred (part of the ‘resurgent’ strategy). Because of a strong spatial component to neural activity in ACC, we trained a separate classifier for left trials and right trials. If there was a trial group—left-bound or right-bound—that had less than ten trials of a trial type—preferred or non-preferred—that accuracy was not calculated. Otherwise, for each of 200 iterations, we used a random 80% of ‘default’ and 80% ‘’ trials to train the classifier. All cells that had a non-zero firing rate were included in the analysis; firing rate for each was calculated over the entire 500 msec decoding window. We performed linear discriminant analysis on the training data set; once the direction normal to the discriminant hyperplane was determined, we selected the best classification threshold from the following three threshold candidates:

- the midpoint between the group centroids
- the intersection point of the probability density functions for the two groups
- the intersection point of the cumulative density functions for the two groups.

Classification accuracy was estimated on the remaining 20% of the dataset, after ensuring that the sample size in each category was the same. For ‘left-side option’ and ‘right-side option’ classifiers, we averaged the accuracy from 200 cross-validation runs for each session. Average accuracy for the better performing of the two classifiers was reporter for each session. Statistics on the estimated classification accuracy was done across sessions.

To control for possible contribution to the difference in ensemble activity between the two conditions from the reward context (high vs low reward), we focused on sessions where reward probability changes were limited to reversals, i.e. where the value of reward probability for each option (‘Left’ and ‘Right’) was either ‘high’ (typically, 0.5 or 0.6) and ‘low’ (typically, 0.25 or 0.3). Because the definition of ‘preferred’ vs ‘non-preferred’ trials is based on the animal’s behavior rather than experimentally defined reward contingencies, and because animals often persisted with preferring one option for some time past the block switch, ‘preferred’ and ‘non-preferred’ trial groups had a mixture of trials that fell within ‘high’ and ‘low’ reward contingency for that choice option. This mixing gave us an opportunity to repeat FR-based classification for trials from the preferred and non-preferred categories that were matched for local reward rates.

To control for possible contribution to the difference in ensemble activity between the two conditions from the way, in which an animal was executing the trials, we also estimated classification accuracy for a linear classifier trained on spatial trajectories rather than ensemble firing rates. The spatial trajectory for each trail was represented either as the mean X and Y positions from 100 msec bins or as the first five principal components of the raw video capture of the animal’s X and Y positions.

## References

Barack, D.L., and Platt, M.L. (2017). Engaging and Exploring: Cortical Circuits for Adaptive Foraging Decisions. In Impulsivity (Springer), pp. 163–199.

Behrens, T.E., Woolrich, M.W., Walton, M.E., and Rushworth, M.F. (2007). Learning the value of information in an uncertain world. Nat Neurosci 10, 1214–1221.

Blanchard, T.C., and Gershman, S.J. (2018). Pure correlates of exploration and exploitation in the human brain. Cognitive, Affective, & Behavioral Neuroscience 18, 117–126.

Blanchard, T.C., and Hayden, B.Y. (2014). Neurons in dorsal anterior cingulate cortex signal postdecisional variables in a foraging task. Journal of Neuroscience 34, 646–655.

Bonini, F., Burle, B., Liégeois-Chauvel, C., Régis, J., Chauvel, P., and Vidal, F. (2014). Action monitoring and medial frontal cortex: leading role of supplementary motor area. Science 343, 888–891.

Brown, J., Behnam, R., Coddington, L., Tervo, D., Martin, K., Proskurin, M., Kuleshova, E., Park, J., Phillips, J., and Bergs, A.C. (2018). Expanding the Optogenetics Toolkit by Topological Inversion of Rhodopsins. Cell.

Brown, J., Pan, W.-X., and Dudman, J.T. (2014). The inhibitory microcircuit of the substantia nigra provides feedback gain control of the basal ganglia output. Elife 3, e02397.

Brown, S.P., and Hestrin, S. (2009). Intracortical circuits of pyramidal neurons reflect their long-range axonal targets. Nature 457, 1133.

Cleland, B.S., Guerin, B., Foster, T., and Temple, W. (2001). Resurgence. The Behavior Analyst 24, 255–260.

Cohen, J.D., McClure, S.M., and Angela, J.Y. (2007). Should I stay or should I go? How the human brain manages the trade-off between exploitation and exploration. Philosophical Transactions of the Royal Society B: Biological Sciences 362, 933–942.

Donoso, M., Collins, A.G., and Koechlin, E. (2014). Foundations of human reasoning in the prefrontal cortex. Science 344, 1481–1486.

Doyon, J., and Benali, H. (2005). Reorganization and plasticity in the adult brain during learning of motor skills. Current opinion in neurobiology 15, 161–167.

Fiser, J., Berkes, P., Orbán, G., and Lengyel, M. (2010). Statistically optimal perception and learning: from behavior to neural representations. Trends in cognitive sciences 14, 119–130.

Gerfen, C., and Paxinos, G. (2004). The rat nervous system.

Hanks, T.D., Kopec, C.D., Brunton, B.W., Duan, C.A., Erlich, J.C., and Brody, C.D. (2015). Distinct relationships of parietal and prefrontal cortices to evidence accumulation. Nature 520, 220.

Hayden, B.Y., Pearson, J.M., and Platt, M.L. (2009). Fictive reward signals in the anterior cingulate cortex. Science 324, 948–950.

Holroyd, C.B., and Coles, M.G. (2002). The neural basis of human error processing: reinforcement learning, dopamine, and the error-related negativity. Psychological review 109, 679.

Karlsson, M.P., Tervo, D.G., and Karpova, A.Y. (2012). Network resets in medial prefrontal cortex mark the onset of behavioral uncertainty. Science 338, 135–139.

Kolling, N., Behrens, T.E., Mars, R.B., and Rushworth, M.F. (2012). Neural mechanisms of foraging. Science 336, 95–98.

Kolling, N., Wittmann, M., and Rushworth, M.F. (2014). Multiple neural mechanisms of decision making and their competition under changing risk pressure. Neuron 81, 1190–1202.

Langen, M., Durston, S., Kas, M.J., van Engeland, H., and Staal, W.G. (2011). The neurobiology of repetitive behavior:… and men. Neuroscience & Biobehavioral Reviews 35, 356–365.

Lee, A.T., Gee, S.M., Vogt, D., Patel, T., Rubenstein, J.L., and Sohal, V.S. (2014). Pyramidal neurons in prefrontal cortex receive subtype-specific forms of excitation and inhibition. Neuron 81, 61–68.

Lu, J., Tucciarone, J., Padilla-Coreano, N., He, M., Gordon, J.A., and Huang, Z.J. (2017). Selective inhibitory control of pyramidal neuron ensembles and cortical subnetworks by chandelier cells. Nature neuroscience 20, 1377.

Ma, L., Hyman, J.M., Durstewitz, D., Phillips, A.G., and Seamans, J.K. (2016). A quantitative analysis of context-dependent remapping of medial frontal cortex neurons and ensembles. Journal of Neuroscience 36, 8258–8272.

McGuire, J.T., Nassar, M.R., Gold, J.I., and Kable, J.W. (2014). Functionally dissociable influences on learning rate in a dynamic environment. Neuron 84, 870–881.

Morishima, M., and Kawaguchi, Y. (2006). Recurrent connection patterns of corticostriatal pyramidal cells in frontal cortex. Journal of Neuroscience 26, 4394–4405.

Naito, A., and Kita, H. (1994). The cortico-nigral projection in the rat: an anterograde tracing study with biotinylated dextran amine. Brain research 637, 317–322.

Nassar, M.R., Wilson, R.C., Heasly, B., and Gold, J.I. (2010). An approximately Bayesian delta-rule model explains the dynamics of belief updating in a changing environment. J Neurosci 30, 12366–12378.

O’Keefe, J., and Recce, M.L. (1993). Phase relationship between hippocampal place units and the EEG theta rhythm. Hippocampus 3, 317–330.

O’Reilly, J.X., Schüffelgen, U., Cuell, S.F., Behrens, T.E., Mars, R.B., and Rushworth, M.F. (2013). Dissociable effects of surprise and model update in parietal and anterior cingulate cortex. Proceedings of the National Academy of Sciences 110, E3660–E3669.

Otchy, T.M., Wolff, S.B., Rhee, J.Y., Pehlevan, C., Kawai, R., Kempf, A., Gobes, S.M., and Ölveczky, B.P. (2015). Acute off-target effects of neural circuit manipulations. Nature 528, 358.

Powell, N.J., and Redish, A.D. (2016). Representational changes of latent strategies in rat medial prefrontal cortex precede changes in behaviour. Nature communications 7, 12830.

Procyk, E., Tanaka, Y., and Joseph, J.-P. (2000). Anterior cingulate activity during routine and non-routine sequential behaviors in macaques. Nature neuroscience 3, 502.

Quilodran, R., Rothe, M., and Procyk, E. (2008). Behavioral shifts and action valuation in the anterior cingulate cortex. Neuron 57, 314–325.

Remondes, M., and Wilson, M.A. (2013). Cingulate-hippocampus coherence and trajectory coding in a sequential choice task. Neuron 80, 1277–1289.

Schuck, N.W., Gaschler, R., Wenke, D., Heinzle, J., Frensch, P.A., Haynes, J.-D., and Reverberi, C. (2015). Medial prefrontal cortex predicts internally driven strategy shifts. Neuron 86, 331–340.

Shenhav, A., Cohen, J.D., and Botvinick, M.M. (2016). Dorsal anterior cingulate cortex and the value of control. Nature neuroscience 19, 1286.

Shenhav, A., Musslick, S., Lieder, F., Kool, W., Griffiths, T.L., Cohen, J.D., and Botvinick, M.M. (2017). Toward a rational and mechanistic account of mental effort. Annual review of neuroscience 40, 99–124.

Stephens, D.W., Kerr, B., and Fernández-Juricic, E. (2004). Impulsiveness without discounting: the ecological rationality hypothesis. Proceedings of the Royal Society of London B: Biological Sciences 271, 2459–2465.

Tervo, D.G.R., Hwang, B.-Y., Viswanathan, S., Gaj, T., Lavzin, M., Ritola, K.D., Lindo, S., Michael, S., Kuleshova, E., and Ojala, D. (2016). A designer AAV variant permits efficient retrograde access to projection neurons. Neuron 92, 372–382.

Vogt, B.A., and Paxinos, G. (2014). Cytoarchitecture of mouse and rat cingulate cortex with human homologies. Brain Structure and Function 219, 185–192.

Youden, W.J. (1950). Index for rating diagnostic tests. Cancer 3, 32–35.

